# Analysis of the recombination landscape of hexaploid bread wheat reveals genes controlling recombination and gene conversion frequency

**DOI:** 10.1101/539684

**Authors:** Laura-Jayne Gardiner, Luzie U. Wingen, Paul Bailey, Ryan Joynson, Thomas Brabbs, Jonathan Wright, James D. Higgins, Neil Hall, Simon Griffiths, Bernardo J. Clavijo, Anthony Hall

**Affiliations:** Earlham Institute, Norwich, NR4 7UZ, UK; IBM Research, Warrington, UK; John Innes Centre, Norwich, NR4 7UH, UK; Department of Genetics and Genome Biology, University of Leicester, Leicester, LE1 7RH, UK; School of Biological Sciences, University of East Anglia, Norwich, NR4 7TJ, UK.

**Keywords:** Wheat, recombination, crossover, gene conversion, QTL

## Abstract

Sequence exchange between homologous chromosomes through crossing over and gene conversion is highly conserved among eukaryotes, contributing to genome stability and genetic diversity. Lack of recombination limits breeding efforts in crops, therefore increasing recombination rates can reduce linkage-drag and generate new genetic combinations. We use computational analysis of 13 recombinant inbred mapping populations to assess crossover and gene conversion frequency in the hexaploid genome of wheat (*Triticum aestivum*). We observe that high frequency crossover sites are shared between populations and that closely related parental founders lead to populations with more similar crossover patterns. We demonstrate that gene conversion is more prevalent and covers more of the genome in wheat than in other plants, making it a critical process in the generation of new haplotypes, particularly in centromeric regions where crossovers are rare. We have identified QTL for altered gene conversion and crossover frequency and confirm functionality for a novel *RecQ* helicase gene that belongs to an ancient clade that is missing in some plant lineages including *Arabidopsis*. This is the first gene to be demonstrated to be involved in gene conversion in wheat. Harnessing the *RecQ* helicase has the potential to break linkage-drag utilizing widespread gene conversions.

## Main Text

There is an evolutionary requirement for genetic diversity across a species. Shuffling of material between homologous chromosomes, or genetic recombination, breaks linkage between genes resulting in offspring that have combinations of alleles that differ from those found in either of the parents. During meiosis, double-strand breaks (DSBs) can generate sequence variation in gametes via the DSB repair model (Szostak et al., 1983). DSBs are resolved either by homologous recombination as crossovers (COs) i.e. the reciprocal exchange of large regions between chromosomes, or otherwise as non-crossovers (NCOs). A minimum of one CO per chromosome during meiosis is a requirement for proper chromosome segregation (Pardo-Manuel De Villena and Sapienza, 2001). When both COs and NCOs are resolved, they can also give rise to gene conversions (GCs) as a mechanism of DSB repair involving the non-reciprocal transfer of short DNA segments between homologous non-sister chromatids (Sun et al., 2012; Halldorsson et al., 2016). GCs can be either allelic, meaning that one allele of the same gene replaces another allele, or ectopic, meaning that one paralogous DNA sequence converts another, they are also involved in the repair of DSBs that occur during mitosis (reviewed in: Chen et al., 2007).

In plant and animal breeding, researchers strive to identify and introduce loci linked to favourable traits ranging from abiotic or biotic stress resistance to agronomic traits. This often also introduces linked undesirable genes and their resulting traits. The goal is to introduce a favourable allele while minimizing linkage drag from surrounding undesirable alleles. Increasing COs at meiosis breaks up linkage groups reducing linkage drag. However, COs are largely absent in centromeric chromosomal regions (Talbert and Henikoff, 2010). GCs contribute to breaking up linkage groups and it has been observed that GCs are prevalent in centromeric regions suggesting that centromeres do experience genetic change but that DSBs in these regions are converted preferentially to GCs (Shi W et al., 2010; Sun et al., 2012). It is therefore important for us to understand recombination and GC if we are to alter their rates to accelerate the induction of novel allelic combinations or to generate stable cultivars. This is particularly important in bread wheat where recombination frequency is low and skewed toward the ends of chromosomes (Darrier et al., 2017). Wheat has a large (16Gb) complex allohexaploid genome and, in the light of recent advances in the wheat’s genomic and genetic resources, it presents an excellent model crop (Brenchley et al., 2012; Clavijo et al., 2017; Krasileva et al., 2017; IWGSC et al 2018.).

In plants over 80 genes have been identified and characterized that are involved in recombination including crossover and gene conversion formation (Mercier et al., 2015; Fernandes et al., 2018; Ziolkowski et al., 2017; Girard et al 2015; Segulea-Arnaud et al., 2015). For example, in *Arabidopsis,* Ziolkowski et al (2017) found that the *HEI*10 meiotic E3 ligase could control crossover recombination. Recombination rates can also be variable; between populations, within a population and even within a chromosome of a single organism (Schnable et al., 1998; Esch et al., 2007). However, in crops such as wheat, little detail is known about the mechanisms that control genome-wide recombination. In wheat, quantitative trait loci (QTL) have been identified that affect recombination frequency, however, there are few QTLs that have been fully characterized to the point of validating candidate genes (Esch et al., 2007; Wingen et al., 2017; Jordan et al., 2018). There is very little known in wheat about the genes or mechanisms controlling GC or GC frequency across its complex genome.

To investigate the recombination landscape in bread wheat we used 13 genotyped recombinant inbred lines (RIL) populations generated from the UK elite variety Paragon crossed with a diverse collection of landraces and elite material (for population details see Supplemental Table S1 and S2). Using the genotypes of the parents we can accurately map CO positions across the genome (Supplemental Figure S1). In addition to large CO blocks, shorter shifts in genotype were frequently encountered across the genome. These have previously been disregarded as potential genotyping errors or issues with local ordering of markers. These short shifts are potential markers for GC events. Recent advances in wheat’s genome sequence assembly and local ordering of contigs into pseudo-chromosomes allows us to more confidently classify shorter shifts as GCs in wheat (Brenchley et al., 2012; Clavijo et al., 2017). Using this methodology, we have characterized: CO and GC frequencies and locations; how different parental crosses effect the recombination landscapes; genomic regions that control CO and GC frequency and identified a *RecQ*-like gene controlling GC frequency in wheat. We have used whole genome sequencing of selected lines to validate our CO and GC calls and to generate a more comprehensive profile of these events across the genome.

## Results

### Analysis of the CO landscape in wheat

For each of the 13 populations, the number of COs per RIL was recorded across the 21 chromosomes (Figure 1a; Supplemental Figure S2; CO-Phenotype, Methods). CO frequencies show a relatively normal distribution independent of the analysed population with outlier RILs observed with high or low CO frequencies in each population. The average CO frequency per RIL remained relatively stable across the 13 populations varying from 40.8 to 51.9 (Supplemental Table S2) consistent with similar studies (Huang et al., 2012; Esch et al., 2007). For each population we calculated the number of RILs sharing an individual CO site. Figure 1b and Supplemental Figure S3 highlight the distribution of shared CO sites; there is a peak of, on average, 10.2% of CO sites that are seen in only one RIL with the remainder shared in 2 or more lines (Supplemental Table S3). The rate of CO conservation steadily declines as the number of RILs increases. The maximum number of RILs with a conserved CO site never exceeds 55.3% of the population size (range 44.6-55.3%). Although we do not expect highly conserved CO locations in populations of this size, the overlap of a proportion of our COs is likely explained by the binning of our SNPs into 20Mbp windows to define COs.

**Figure 1.**
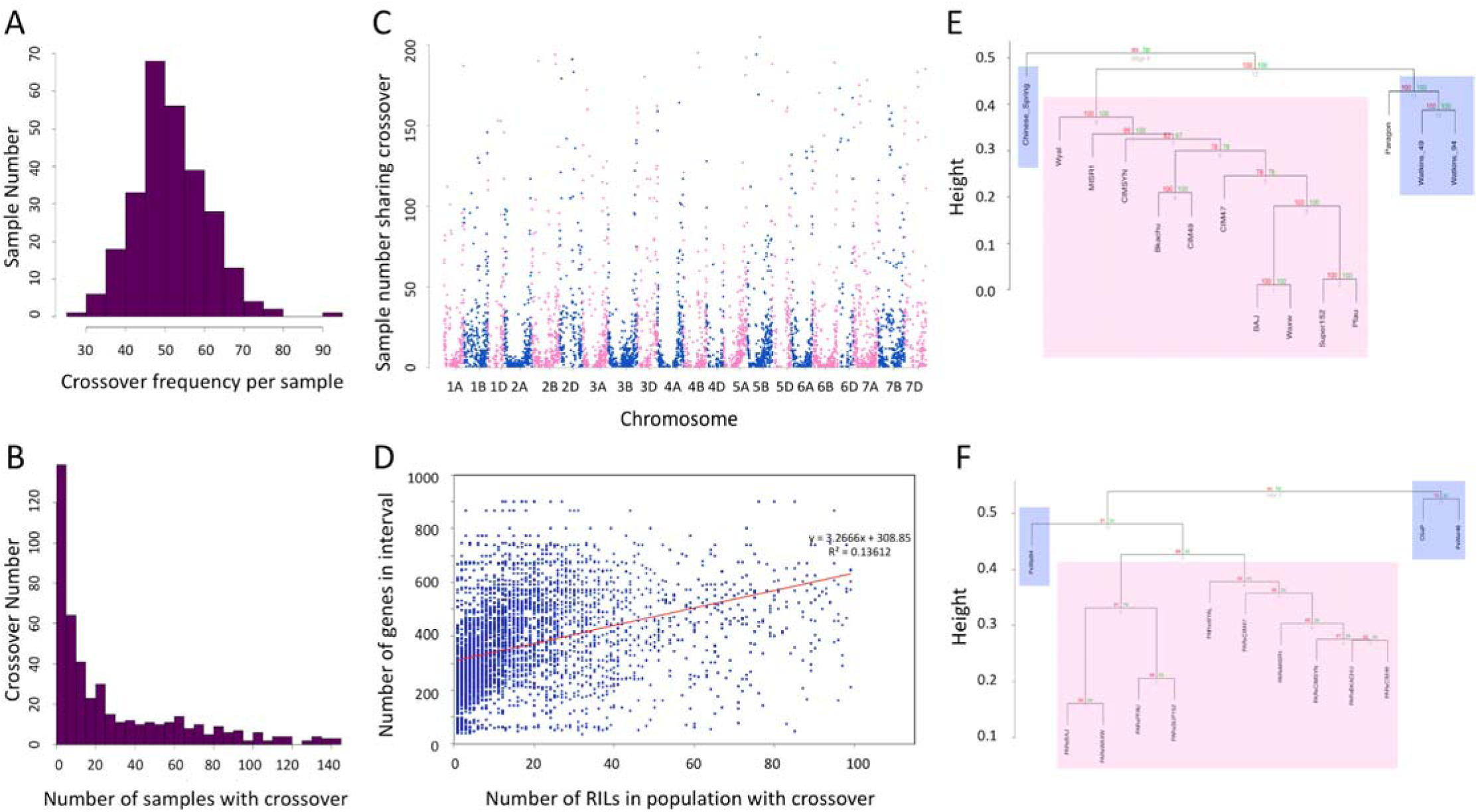
Recombination landscape of wheat. (A) The number of COs recorded for each RIL in the Paragon x Chinese Spring population (CO frequency per sample) as a frequency histogram. (B) **T**he number of RILs sharing each recorded CO (Number of samples with each CO) as a frequency histogram for the Paragon x Chinese Spring population. (C) For all analysed COs the location of the CO (the start of the window that shows a different predominant parental allele compared to the previous window) is plotted on the x axis with the number of samples in the population that share the CO on the y axis. (D) The intersection of two 20Mbp windows defines a CO. Therefore, for all windows of 40Mbp encompassing a central defined CO, the number of high confidence genes that are found within each interval are plotted alongside the number of RILs within each population showing the CO. (E) Parental founders for the 13 populations clustered according to their representative alleles from the 35K SNP array. (F) The 13 populations clustered according to their individual CO profiles i.e. number of RILs with each recorded CO in the population. The dendrograms in (E) and (F) were produced using the R package pvclust average linkage method with correlation-based dissimilarity matrix and the value of this distance metric between clusters is represented as height on the y-axis. AU (Approximately Unbiased) p-values were computed by multiscale bootstrap resampling (bootstrap number of 1,000). Landraces are highlighted with blue boxes and pure breeding lines are highlighted with pink boxes.

We observed that CO sites cluster towards the end of chromosomes and that this effect is more pronounced if the CO site is more frequently encountered i.e. appears in multiple RILs (Figure 1c). CO sites show a bias to genic regions that is statistically significant (two-tailed t test p<0.0001, t = 6.2534, df = 10174). In the regions defined containing COs, the average number of genes is 363.83 and the average proportion of the interval represented by genes is 2.65% whereas across all array SNP intervals the average number of genes per interval decreases to 342.98 and the average proportion of each interval represented by genes decreases to 2.54% (Methods). Further to this bias of COs to genic regions, there was a significant positive correlation between the number of genes at the CO site and the number of RILs with the CO (Figure 1d, Pearson correlation coefficient (r)= 0.271706, n=4780, p<0.00001). Therefore, if a CO is observed in more RILs, then it is more likely to be gene associated, perhaps yielding a favourable phenotype.

We assessed the number of RILs that each identified CO was observed in and found that COs that are more frequently seen within a population are more likely to be seen in all 13 populations. The number of COs that are seen in only one population decreases as more RILs in the population have the CO, conversely COs that are seen in all 13 populations increase in number as more RILs within the populations have the CO (Supplemental Figure S4; Supplemental Table S4).

We clustered parental founder accessions for all of the populations according to their SNP profiles (Figure 1e) and clustered the resultant populations based on their CO profiles (Figure 1f). The CIMMYT lines (pink in Figure 1e and f), cluster together closely and are distinct from the landraces (blue). This is true for the genotype profiles of the parental founders as well as for the resultant CO profiles of the RIL populations. Additionally, from Figure 1e we were able to define pairs of the most closely related parental lines; Baj/Waxw, Bkachu/CIM49 and Super152/Pfau. Looking at the resultant populations after these parents were crossed with Paragon, these are also the most closely related populations according to CO profile. We conclude that more similar parental founders for a population lead to populations with more similar CO profiles. Therefore, wide crosses not only bring in novel alleles but also new COs, potentially breaking linkage groups.

### GCs are more prevalent than COs in wheat

Previously we defined COs by comparing windows that were representative of cM bins (CO-Phenotype, Methods). This methodology is unaffected by subtle differences in genome organization between wheat accessions, such as GC events. Sun et al. (2012) report 120-222 DSBs per meiosis in *Arabidopsis* and although associated GCs should be possible for all of the DSBs, they predict a GC rate of 60-111 assuming that mismatch repair restores 50% of DSBs to their original allelic state. With these rates of GC frequency and previous approximations of GCs being between 2bp-10Kbp in length (Yang et al., 2012; Sun et al., 2012), it is likely that such events will be missed by our analysis with on average 4335 SNPs available for analysis per population. However, after adjustment for our parallel definition of COs, we were able to identify on average 104 potential GCs per RIL across the 13 populations (Supplemental Table S2; Figure 2a; Supplemental Figure S5; GC-Phenotype, Methods).

**Figure 2.**
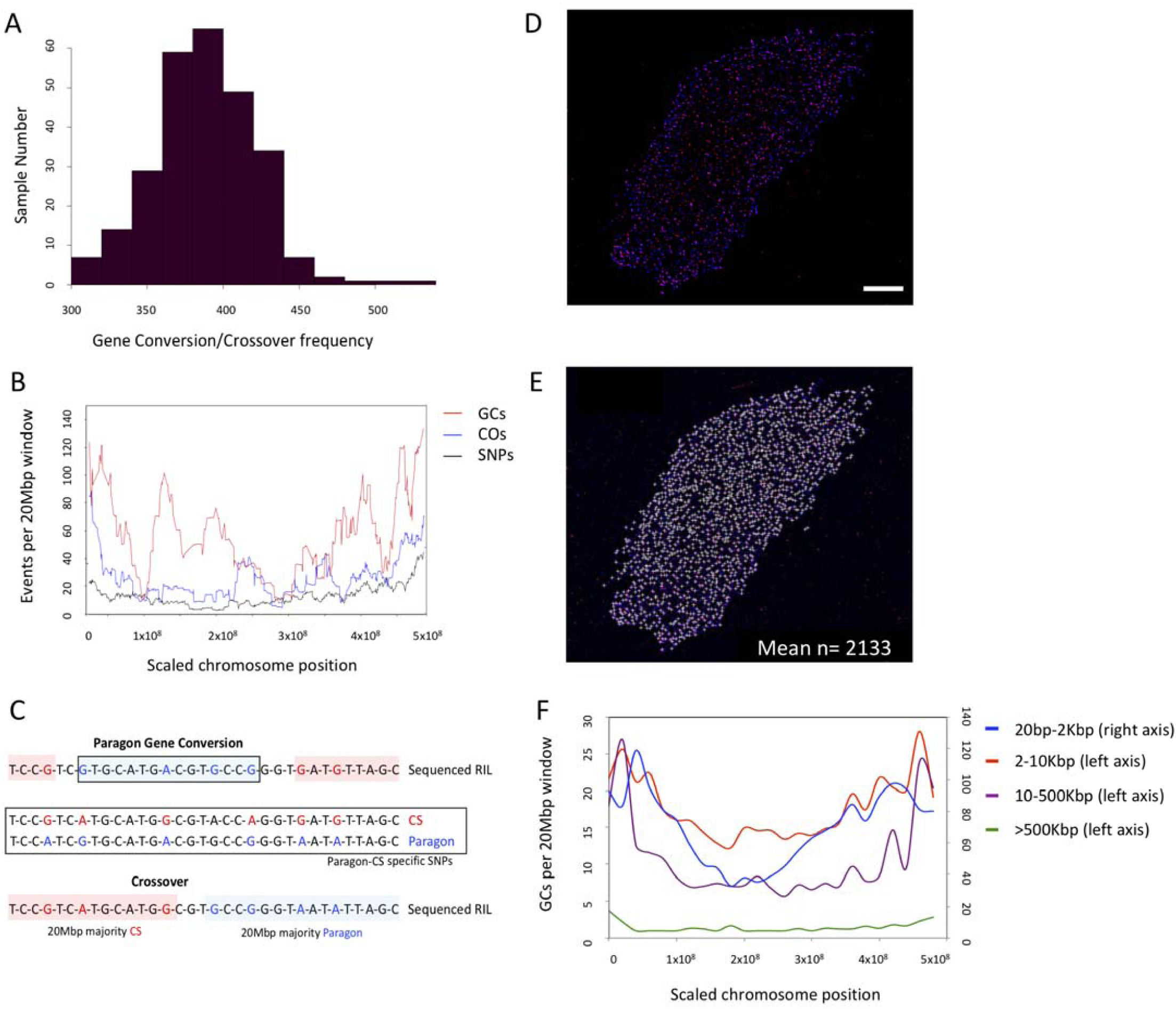
Fine scale analysis of sequence exchange events. (A) The number of COs and/or GCs recorded for each RIL in the Paragon x Chinese Spring population (GC/CO frequency per sample) as a frequency histogram. (B) Line plots separately for; the number of COs (COs), GCs (GCs) and array SNPs per 20Mbp window across the genome. All chromosomes are normalized to 500Mbp in length to be displayed in a single plot. The moving average of each dataset is displayed (period=15). (C) Schematic of methodology for calling Gene Conversions (GCs) and Crossovers (COs) in the skim sequencing data using pre-defined Paragon and Chinese Spring specific homozygous SNPs. (D) Immunolocalization of the chromosome axis protein ASY1 (blue) and yH2A.X (red) a marker for DNA DSB on hexaploid wheat leptotene male meiotic nuclei. Scale bar = 10 μM. (E) Original nuclei as per (D) however, yH2A.X foci are marked that co-localise with ASY1. Mean number of yH2A.X foci across 5 replicates shown, from displayed image n=1673. (F) Line plots separately for; the number of GCs 20bp-2Kbp, 2-10Kbp, 10-500Kbp and >500Kbp in length per 20Mbp window across the genome. Chromosomes are normalized as per (B) and the average frequency per window is displayed.

In Figure 2b we compare the distribution of COs to the profile of GCs across the genome. Both distributions show the characteristic increase in frequency towards the distal regions of the chromosomes indicative of our likely detection of GC events associated with COs and therefore observed at similar locations. However, high GC frequency in the centromeric regions, where COs are sparse, was also seen and is likely to represent GCs associated with NCOs.

### Using whole genome sequencing to define GC locations

There are three hypotheses as to how we are able to identify potential GCs in RILs using low resolution genotyping; either gene conversion events are far more prevalent in wheat than *Arabidopsis* and other eukaryotes, lengths of GCs can be longer in wheat or the GCs could represent genotyping error or structural variation in the RILs. To test these hypotheses and more precisely map COs and GCs we performed whole genome sequencing at low coverage (skim sequencing) for 12 lines from the Paragon x Chinese Spring population. These lines represent the upper and lower ends of the CO and GC frequency range (Supplemental Table S5). We defined 31,327,143 homozygous SNPs between Paragon and Chinese Spring along the Chinese Spring IWGSC RefSeq V1.0 reference sequence (IWGSC et al., 2018) that represent parent specific allelic differences for the RIL population (Methods). This translates to 1 SNP approximately every 540bp to discriminate Chinese Spring and Paragon that is the resolution of our event detection. Skim sequencing data for each of the 12 lines was aligned to RefSeq V1.0 gaining on average 6.42X coverage across 91.3% of the genome and the defined parental SNP set was used to identify homozygous Paragon and Chinese Spring specific alleles within the sequencing data to allow us to identify COs and GCs (Figure 2c; Methods).

The high-resolution whole genome skim sequencing allowed us to identify on average 30,110 potential GC events per RIL. We adapted categories used by Yang et al (2012) to assign confidence to our skim sequencing GCs by classifying them according to length (Methods; Table 2). We noted that GC numbers could be inflated as an artefact of incomplete or locally inaccurate reference genome assemblies or due to structural variation between the analyzed lines and the reference genome. Structural variants are likely to present as events that are conserved across the analyzed lines. We used the abundance levels of our events across the analyzed lines to calculate a false positive rate for our calls with regard to structural variation. 49.3% of our identified events were unique to a single RIL with 82.4% observed in <50% of the analysed lines. We can assign higher confidence to our unique GCs. Interestingly; the average length of unique GCs was 452Kbp. However, for those GCs conserved across the majority of lines (>50%) the average length was only 3.7Kbp. Therefore it appears that the shorter GCs that we identified are more likely to represent structural variation between Chinese Spring and Paragon. Considering only our high confidence GC calls or unique calls we defined a range of 7,909-22,847 and on average 14,795 GCs events per RIL, of which, on average 10,064 GCs were of 2bp-2Kbp, 2,255 GCs of 2-10Kbp, 2,247 of 10-500Kbp and 228 of >500Kbp in length (Table 2, Methods). As a final validation we identified sequencing read pairs that spanned our GC shifts from Chinese Spring to Paragon encompassing both a Chinese Spring and a Paragon SNP (Methods). This analysis provided an accuracy rate of 85.2% for our definition of GC shifts. The false positives (14.8%) are likely to be a result of read misalignment or structural reference related errors.

**Table 2.**
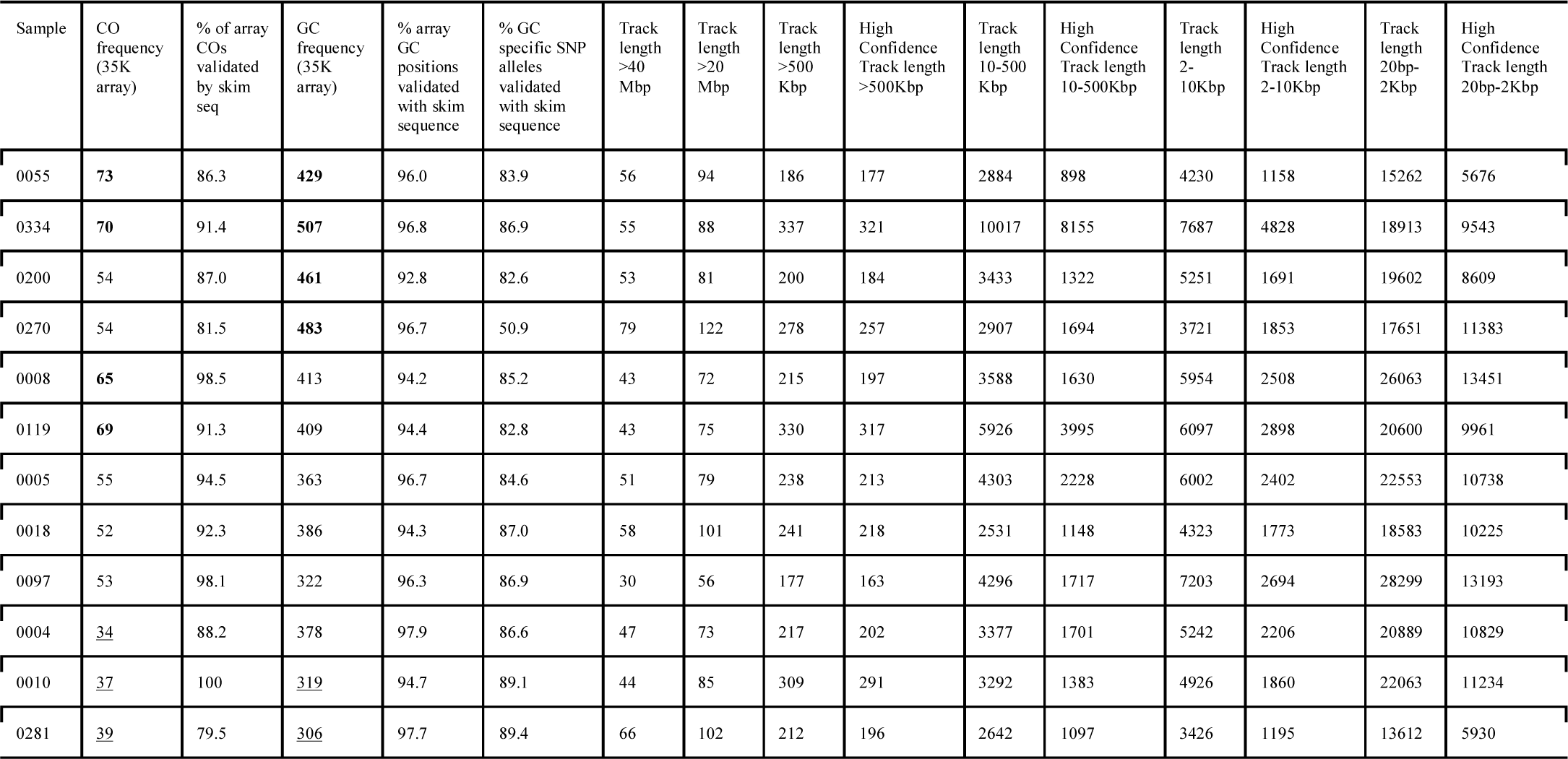
Sequence exchange events defined from whole genome skim sequencing. Detailing for the 12 Paragon x Chinese Spring samples skim sequenced; the sample number, CO and frequency defined using the 35K array, % of these events validated by skim sequencing, GCs identified with track lengths >40,000,000bp (representative of COs), >20,000,000bp, >500,000bp, 10,000bp-500,000bp, 2,000bp-10,000bp and 20bp-2,000bp. High confidence GCs are defined as those where the specific array SNP allele that was used to make the call could be validated in the skim sequencing.

In *Arabidopsis*, Sun et al. (2012) predict GC rate based on the DSB frequency per meiosis. Here, to allow us to perform a similar validation for our GC calls, we calculated the DSB rate for male hexaploid wheat leptotene nuclei (n=5) using immunolocalization recording on average 2133+/-157 DSBs per meiosis (Figure 2d and 2e; Supplemental Figure S6). This translates to 3952-4580 considering female DSBs per meiosis, and when considering that we are observing an F8 generation we would therefore expect 31,616-36,640 DSBs to have occurred in our analyzed RILs. Although, since our F8 RILs have passed through single seed descent becoming increasingly homozygous in each resultant generation, detectable allelic GCs, where an allele of the same gene replaces another variant allele, will decrease as generations increase. Accounting for increasing homozygosity at a rate of approximately 50% per generation, we estimate that 7874-9125 DSBs would have the potential to be detectable allelic GCs. It is unclear how DSBs directly translate to detectable GCs in wheat and the impact of mismatch repair on this number however, when we focus on our high confidence GC calls from the skim sequencing, after correction for our classification accuracy rate of 85.2% we define a range of 6,588-19,192 and on average 12,411 GCs of 20bp-500Kbp that overlaps the frequency of DSBs.

We showed with array SNPs that GCs increase in frequency towards the distal regions of the chromosomes but also show higher frequency in centromeric regions. Here we profiled the different length high confidence GCs defined from the whole genome sequencing to see if their profiles differ (Figure 2f). It is evident from Figure 2f that all GCs tend to increase at the distal regions of the chromosomes, however, compared to GCs of >500Kbp in length, which are likely to include crossovers, both 10-500Kbp and 2-10Kbp GCs maintain an elevated frequency that is conserved across proximal and centromeric regions. Shorter GCs of 20bp-2Kbp display the highest frequency that is conserved highly across proximal regions and to a relatively high level at the centromere. It is therefore the GCs from 20bp-500Kbp in length, which appear to largely break proximal and centromeric regions.

### Using whole genome sequencing to validate our array based CO and GCs

Across the 12 sequenced RILs we observed 52 COs per RIL considering events of at least 40Mbp which closely reflects the average of 54.6 COs for the same 12 RILs defined using the array SNPs. This is likely because the 40Mbp represents the total analysis window that we used for array SNPs to define a central CO at the intersection of two 20Mbp windows. Furthermore, Table 2 highlights that we could validate up to 100% of the COs from the array analysis with the skim sequencing datasets and that they overlapped events that averaged 42,859,250bp in length with 84.7% of events >500Kbp. On average we validate 90.7% of COs with the skim sequencing data.

Similarly, we validated GC calls from the array analysis using the skim sequencing. We could directly associate on average 95.7% of our previously defined GCs with GC events in the sequencing data that were on average 29,992,400bp in length with 61.2% of events <20Mbp (Table 2). Furthermore on average we could validate 83.0% of the specific array SNP alleles that were used to define GCs and given the reported agreement rates of 85.7% for SNP arrays when compared to sequencing data by Burridge et al. (2017), this is in line with expectations. Using the array, the reason for our ability to identify GCs in low-resolution genotyping data is that we typically detect the larger GC events whereas whole genome sequencing gives the ability to robustly detect shorter GC events across the genome. It appears the frequency of GCs is far higher than in *Arabidopsis* and in contrast to other eukaryotes studied to date, longer GCs are also prevalent in wheat (Sun et al., 2012).

### QTLs identified for CO frequency

In our analysed RIL populations we observed a normal distribution of CO and GC frequency with a positive skew (Figure 1a and 2a). As such, there were subsets of RILs with increased levels of GCs or COs compared to the population average. Therefore, we used GC and CO frequency as traits for QTL analysis (Methods). The CO analysis (CO-Phenotype, Methods) identified a robust QTL for the Paragon x Chinese Spring population that explained >6% of the variation (LOD score 3.64, p<0.05; Table 1; Figure 3a, 3c and 3e). The Holliday junction ATP-dependent DNA helicase *RuvB*-like was located within ∼1.5 Mbp of the QTL peak on chromosome 6A. This gene is known to act in a complex with *RuvA* to promote strand exchange reactions in homologous recombination (Shalev et al., 1999) and is therefore our main candidate for the QTL. The *RuvA-*like gene was also located within the QTL interval (∼4 Mbp from main peak).

**Table 1.** QTLs identified for CO frequency. Detailing for the QTL analysis; the population under analysis, phenotype, cross type and generation, chromosome, position (cM), logarithm of odds (LOD) score, % variation explained, additive effect, flanking markers including QTL region length and gene number, top marker from QTL scan and annotation of identified candidate genes in the region.

| Population/<br>Phenotype | Cross | BC.<br>gen | F.gen | Ch<br>r | Pos<br>cM | LOD | %var | Add eff | Flanking markers | Top marker | Candidate genes plus annotation | Parent<br>with<br>increasing<br>effect |
| --- | --- | --- | --- | --- | --- | --- | --- | --- | --- | --- | --- | --- |
| CSxP<br>CO-Phenotype | RIsself | NA | NA | 6A | 11.84 | 3.637 | 6.037 | -4.502 | AX-95155001<br>AX-94507146<br>6,740,223-21,438,976bp<br>TraesCS6A01G013800-TraesCS6A01G040900<br>(including 267 genes) | AX-94901219<br>11.84cM<br>6D: 13,554,815 bp;<br>TraesCS6D01G032100<br>; Pre-mRNA-processing protein 40A<br>Marker at 11.84 cM on<br>6A: 14,609,435bp | **TraesCS6A01G026300.2 Holliday**<br>**junction ATP-dependent DNA helicase**<br>**RuvB (13,063,092 bp)**<br>TraesCS6A01G037500.1 Holliday<br>junction ATP-dependent DNA helicase<br>RuvA (18,534,777bp) | Chinese<br>Spring |
| CSxP<br>GC-Phenotype | RIsself | NA | NA | 2A | 161 | 2.513 | 4.212 | -14.73 | AX-94616756<br>AX-94639168<br>51,940,072-70,183,206bp<br>TraesCS2A01G098600-TraesCS2A01G119800<br>(including 207 genes) | AX-94419084<br>57,999,746 bp | **TraesCS2A01G103300 (56,942,731 bp)**<br>**ATP-dependent RNA helicase Dead;**<br>**DNA helicase, ATP-dependent, RecQ**<br>**type** | Chinese<br>Spring |
| PxCIM47<br>GC-Phenotype | RIsself | 0 | 5 | 2B | 0 | 6.041 | 6.623 | 0.519 | AX-94768203<br>AX-94546045<br>47,425,660-64,988,255bp<br>TraesCS2B01G084900-TraesCS2B01G104200<br>(including 189 genes) | AX-94768203<br>64,988,255 bp | **TraesCS2B01G103300 (64,091,671bp)**<br>**ATP-dependent DNA helicase PIF2** | CIM47 |
| PxWat94<br>GC-Phenotype | RIsself | 0 | 4 | 4B | 191 | 2.591 | 10.461 | -0.838 | AX-94840592<br>AX-95186017<br>3,513,381-4,996,179 bp<br>TraesCS4B01G005000-TraesCS4B01G007500<br>(including 26 genes) | AX-94840592<br>4,996,179 bp | **TraesCS4B01G006700 Protein HIRA**<br>**(4,526,708bp)** | Paragon |
| PxBaj<br>GC-Phenotype | RIsself | 0 | 4 | 5A | 304.6 | 3.802 | 9.898 | 1.24 | AX-95240942<br>AX-94480416<br>464,478,676-487,684,559bp<br>TraesCS5A01G249800-TraesCS5A01G277700<br>(including 273 genes) | AX-94758045<br>472,344,060bp | **TraesCS5A01G257000**<br>**(472,343,789-472,347,557bp)**<br>**WPP domain-interacting protein 1**<br>TraesCS5A01G260500 (474,537,170bp)<br>Regulator of chromosome condensation<br>(RCC1)<br>TraesCS5A01G381700LC (472,221,190-472,221,614bp)<br>BURP domain protein RD22-similar to C-PHI for homologous chromosome pairing | Baj |

**Figure 3.**
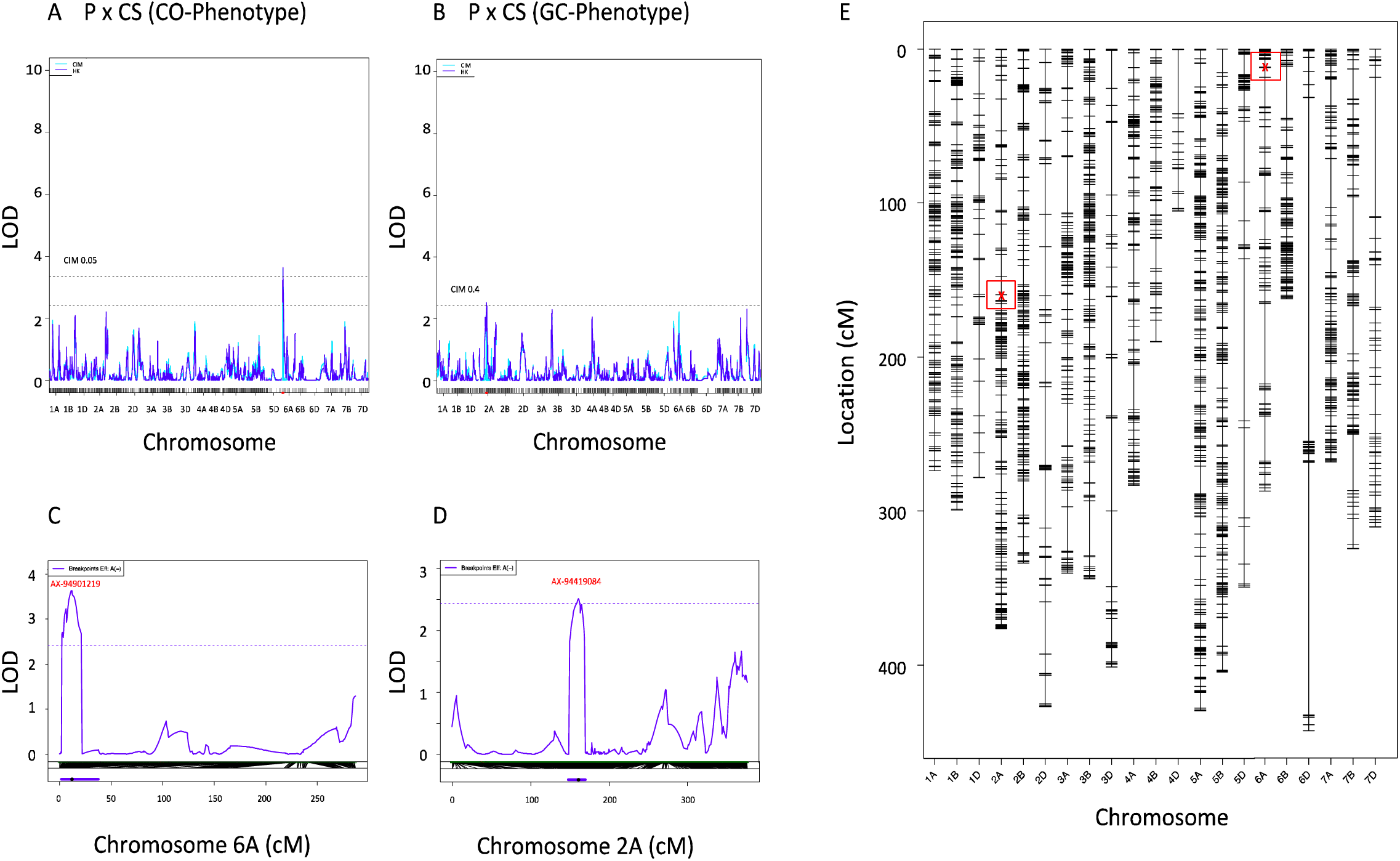
Output from QTL analysis from the Paragon x Chinese Spring population. QTL analysis output for the Paragon x Chinese Spring population that yielded significant associations for either **(A)** CO-Phenotype or **(B)** GC-Phenotype (p<0.05). Detailing LOD scores plotted over the respective linkage groups i.e. chromosomes. Increased resolution of QTL peaks for **(C)** CO-Phenotype and **(D)** GC-Phenotype. Finally, **(E)** shows the locations of the array SNPs showing the peak associations marked in red surrounded by a red box while also showing all other array SNP locations per chromosome.

For the GC frequency trait (GC-Phenotype, Methods) we identified multiple robust QTL that explained 4.2-10.5% of the observed variation (Table 1; Figure 3b, 3d and 3e; Supplemental Figure S7). We only identified QTL for four of the 13 populations likely due to the low power of our analysis in some of the populations since they were made up of <100 RILs. We studied genes within the QTL intervals and identified four gene candidates for GC frequency that were on average 600Kbp from the QTL peak and showed functional significance (p<0.05); firstly, ATP-dependent RNA helicase *RecQ-like* (for the purpose of this paper we have named this *RecQ-7*) on chromosome 2A from the Paragon x Chinese Spring population analysis, overexpression of *RecQ* in rice embryogenic cells has been linked to stimulation of homologous recombination (Li et al., 2004) (Figure 3b); secondly, ATP-dependent DNA helicase *PIF2* on chromosome 2B from the Paragon x CIMMYT 47 analysis, *PIF2* is a known DNA repair and recombination helicase (Supplemental Figure S7a); thirdly, a gene encoding the protein HIRA on chromosome 4B from the Paragon x Watkins 94 analysis, chromatin reassembly during DSB repair has been shown to be dependent on the HIRA histone chaperone (Li, X. and Tyler, J. 2016) (Supplemental Figure S7b); finally, WPP domain-interacting protein 1 on chromosome 5A from the Paragon x Baj analysis, this gene is key for nuclear assembly and transport and is involved in the same pathway as the gene *RCC1* that is seen in the same interval (Supplemental Figure S7c). We noted a low confidence gene ∼120Kbp from the WPP domain-interacting protein 1 in the Paragon x Baj analysis, this BURP domain protein RD22 shows similarity to the gene *C-Ph1* that is involved in homologous chromosome pairing and could also be contributing to this QTL peak (Griffiths et al. 2006).

### Validation of candidate genes

We used the Cadenza TILLING population, a mutagenized bread wheat population that has been widely characterized using exome sequencing, to identify lines with likely knockouts of our candidate genes from the QTL analysis, we prioritized stop codon inducing mutations that were as close to the start of the gene as possible to ensure a null phenotype (Krasileva et al. 2017) (Supplemental Table S6 and S7). We then use the high frequency background EMS mutations in the TILLING lines to call COs and GCs as previously, however, here recording homozygous/heterozygous shifts across the genome (Methods) (Wijnker *et al.,* 2013). Firstly, for the Holliday junction ATP-dependent DNA helicase *RuvB-like* that was associated with CO frequency in the Paragon x Chinese Spring population, the CO frequencies of the eight knockout *RuvB* lines and the control group of ten lines showed no discernible difference with average frequencies of 57.4 and 57.6 respectively (Figure 4a; Supplemental Table S6; Supplemental Note S1; Methods).

**Figure 4.**
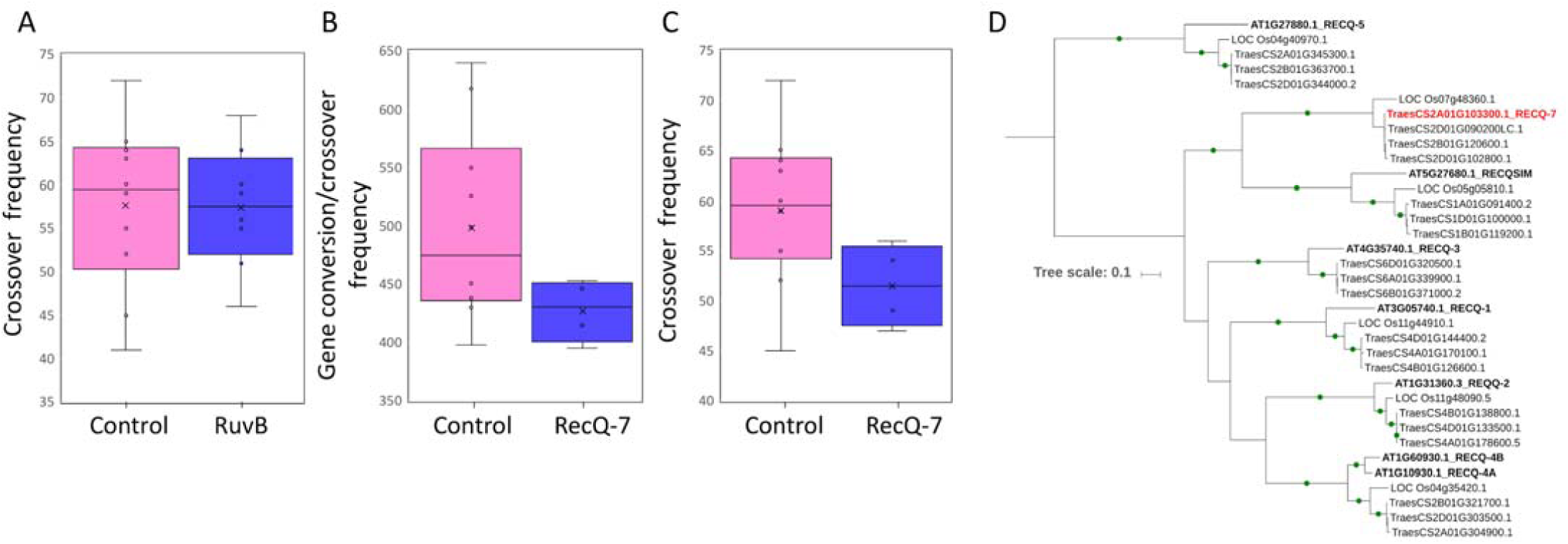
Examination of candidate genes from QTL analysis *RecQ-7* and *RuvB*. (A) Box plot comparison of the knockout *RuvB* lines with the control lines, defining CO frequency using collapsed linkage windows as per CO-Phenotype. (B) Box plot comparison of the knockout *RecQ-7* lines with the control lines, defining CO/GC frequency using GC-Phenotype. (C) Box plot comparison of the knockout *RecQ-7* lines with the control lines, defining CO frequency using CO-Phenotype. (D) Phylogenetic tree of identified genes across multiple species (*Arabidopsis,* rice and wheat) with sequence similarity to the *RecQ* helicase family, including our wheat candidate *RecQ-7* gene for comparison. Bootstrap values >= 90 % are shown as green dots on the branches.

Secondly, for the ATP-dependent RNA helicase *RecQ-7* that was associated with GC frequency in the Paragon x Chinese Spring population, we observed a significant decrease in GC frequency in the *RecQ-7* mutant knockout group of four lines compared to the control group (Welch t test, P=0.0338, t=2.424, df=11) (Supplemental Note S2). We observed average GC frequencies of 426.96 and 498.54 for mutant and control lines respectively (Figure 4b; Supplemental Table S6; Supplemental Note S3; Methods). We then went on to define CO frequency for the *RecQ-7* knockouts to determine if our GC-Phenotype QTL, which mainly reflects increased GC, could translate to an increase in COs. We observed a significant decrease in CO frequency in the *RecQ-7* mutant group compared to the control group (Welch t test, P=0.0411, t=2.3439, df=10) with average CO frequencies of 51.5 and 59 for mutant and control respectively (Figure 4c, Methods).

In *Arabidopsis* there are 7 *RecQ* genes, *AtRecQl 1, 2, 3, 4A, 4B, 5* and *AtRecQsim*. *RecQ4A* and *RecQ4B* have been identified as playing a role in recombination (Hartung et al. 2007; Séguéla-Arnaud et al. 2015). To understand how the wheat *RecQ7* gene is related to the *Arabidopsis RecQ* genes we built a phylogenetic tree including rice, *Arabidopsis* and wheat *RecQ* homologs (Figure 4d). We found that the *RecQ7* gene failed to cluster with any of the *Arabidopsis* genes, instead it clusters in a very well defined separate branch with the rice *RecQ-Like* gene Os07g48360 defining a novel *RecQ* clade that has been lost in some plant lineages (Supplemental Figure S8; Hartung et al. 2006). We used the Cadenza TILLING population to ascertain if knockouts of the three homoeologous wheat homologs of the *Arabidopsis* candidates *RecQ4A* and *RecQ4B,* identified from Figure 4d, showed similar CO phenotypes (Supplemental Table S6). Knockouts of our homoeologous gene candidates resulted in general decreases in the average CO frequency per line compared to the control group, however none were statistically significant (Supplemental Note S4).

We next used the Cadenza TILLING population to ascertain if knockouts of the homoeologous gene copies of our candidate gene *RecQ-7* on chromosome 2A, showed similar GC phenotypes to *RecQ-7* (Supplemental Table S6; Supplemental Note S5; Supplemental Figure S9). Knockouts of our homoeologous gene candidates resulted in a decrease in the average GC frequency per line from 498.5 in the control group to 414.4 and 449.4 for sub-genomes B and D respectively (GC-Phenotype). When we consider all sub-genome A, B and D knockouts as a group together there is a significant decrease in GC frequency between this group and the control group (Two tailed t test, P=0.0371, t=2.2617, df=17).

The increasing effect of *RecQ-7* on CO frequency was associated with the landrace parental line Chinese Spring. We compared our candidate gene sequence between Paragon and Chinese Spring using genomic and transcriptome sequence and saw a high level of sequence conservation, however, there was an insertion of 12 bp in Paragon compared to Chinese Spring that is potentially a microsatellite expansion in the intronic sequence. Furthermore, Cadenza, the founder accession for the TILLING population that we used to knockout the gene and observe a phenotypic effect, has a homologous *RecQ-7* sequence that is more comparable to the genic sequence seen in Chinese Spring. In Chinese Spring, *RecQ-7* is reportedly expressed in a large range of tissues but notably it is expressed at the highest levels in the vegetative and reproductive spike at anthesis, flag leaf stage, ear emergence and meiosis (Borrill *et al.*, 2016). In the vegetative spike, our candidate homoeolog *RecQ-7* on sub-genome A is expressed at 1.4X and 1.7X the level of the homoeologs on sub-genome B and D respectively. Similarly in the reproductive spike *RecQ-7* on sub-genome A is expressed at 1.3X and 1.9X the level of the homoeologs on sub-genome B and D respectively. Therefore, although a combined triple gene knockout may be beneficial, this could be why a significant observable effect is only gained from the knockout of our single candidate *RecQ-7* on sub-genome A with uneven gene expression across the *RecQ-7* homeologs that is biased to sub-genome A. However, it may be that a triple mutant would be lethal.

We focused on genes that were identified from the Chinese Spring x Paragon population since this was largest population with the highest number of SNPs available for analysis and was therefore likely to give the most robust candidates. Knockouts of QTL candidates from the other populations showed observable decreases in GC/CO frequency, however they showed decreases that were typically not statistically significant (Supplemental Note S6; Supplemental Table S7). The notable exception was the WPP domain-interacting protein 1 from the Paragon x Baj population analysis where, although mutants for this gene showed a decrease in GC frequency that was not significant, this translated to a significant decrease in in CO frequency in the mutant group compared to the control group. This gene could be an important candidate for further analysis. It is possible that combined triple gene knockouts for our additional QTL candidates may yield significant results.

## Discussion

Using the 35K array we see the characteristic normal distribution of CO frequency, with an average of 41-52 COs per RIL, and a non-random distribution of COs between RILs and along the chromosomes (Huang et al., 2012; Esch et al., 2007; Darrier et al., 2017). There is a positive correlation between the number of genes in the CO interval and the number of RILs with the CO suggesting that perhaps recombination in gene rich regions leads to favourable phenotypes that are under selective pressure. In addition, we show that introducing a more divergent parental founder e.g. a landrace such as Chinese Spring or a Watkins accession to be crossed with an elite line could increase the potential for introducing new CO profiles in the resultant population in addition to new allelic diversity.

Using whole genome skim sequencing we profile GC events across the wheat genome revealing a large unexplored source of sequence exchange between chromosomes that is particularly desirable in the drive to break linkage drag in the crossover sparse centromeric regions. We show that larger GC events can be profiled using low-density genotyping arrays where previously the crop breeding and research community may have ignored such changes regarding them as potential genotyping errors. These large GCs have not been described before in eukaryotes and may be the result of large complex repetitive polyploid genomes. There is also the possibility that these large GCs could actually be double COs where *RecQ-7* could promote more intermediates going down the non-interfering CO pathway. However, we do not observe a global increase in non-interfering COs, although we cannot rule out a possible role in recombination intermediate branch migration by *RecQ-7* as observed in the human ortholog BLM (Karow et al., 2000), that may separate components of a double Holliday junction substantially enough for both Holliday junctions to resolve via the class II pathway as COs. In any case, this would be an equally useful tool to reduce linkage drag and an interesting feature of such a complex polyploid.

Despite the critical role of GC in genome diversity and evolution little is known about the mechanism of control. In yeast the protein complex MutLβ and meiosis-specific helicase, *Mer3* have been implicated controlling the length of gene conversions (Duroc et al., 2017). Here, we were able to identify multiple QTL affecting GC frequency. We have identified a candidate helicase gene, ATP-dependent RNA helicase *RecQ*, on chromosome 2A that we named *RecQ-7.* Using EMS induced gene knockouts, we went on to validate our gene candidate *RecQ-7* observing a drop in GC frequency in those lines where our candidate gene was knocked out. This reduction in GC frequency also correlated with a significant decrease in CO frequency. Furthermore, the decrease in GC was observed when either of the other homoeologous gene copies was knocked out. By performing phylogenetic analysis we determined that *RecQ-7* was present in rice but not in *Arabidopsis*. Therefore, genetic screens to identify suppressors of recombination in *Arabidopsis* would not identify *RecQ-7*, although *RecQ-4* might perform a similar role (Seguela-Arnaud et al., 2015). However, *RecQ-7* (OsRecQ 886 in rice) is structurally divergent from *RecQ-4* (reviewed in Hartung and Puchta, 2006). *RecQ-7* is structurally similar to moss *RecQ-6* that is essential for gene targeting, but has no apparent role in DNA repair (Wiedemann et al., 2018). Therefore *RecQ-7* is an important target for wheat breeders to increase recombination in genomic ‘cold’ regions by increasing CO and GC frequency and could be transferred to dicotyledonous crops to be used as an enhancer of recombination.

## Materials and Methods

### Crossover calls (CO-Phenotype)

The SNP sequences from the 35K array were aligned (anchored) to the IWGSC RefSeq V1.0 wheat genome reference sequence (IWGSC et al., 2018) and therefore, for each SNP, we have its exact base pair position the Chinese Spring reference. To identify if specific sequences or regions are targeted for recombination consistently between populations, the SNP Chinese Spring reference anchoring was implemented for all populations throughout this study to aid comparative analyses between populations. For each population, for each RIL, array SNPs were annotated according to the parental allele that they represented. SNPs that were deemed to be so close together as to render recombination between them more unlikely were collapsed into bins representative of cM bins. With a genome size of 16Gb and a previously reported cM genome size of 3894cM we collapsed SNPs within intervals across the genome of 20Mbp (16Gbp/3894cM * 5 the estimated shared haplotype length around a SNP in wheat) (Huang et al., 2012). Collapsing SNPs involved classifying the most frequently encountered parent specific allele per window, if the window appeared to be more Parent 1 or 2 i.e. homozygous for Parent 1 or 2 this was recorded, and a mixture of the two parents was recorded as a heterozygous region. A change in the most frequently encountered parent specific allele from one window to the next across the genome was recorded as a CO. This methodology is unaffected by subtle differences in genome organization between wheat accessions and is therefore a conservative method for calling COs enabling cross-population comparisons, our refinement of a CO will be limited to the intersection of two 20Mbp windows.

To call COs from the Cadenza TILLING population, for each of the lines we assessed, we utilized SNPs that were called by aligning exome capture data to the IWGSC RefSeq V1.0 Chinese Spring reference sequence. The SNPs are publicly available at (https://plants.ensembl.org/Triticum_aestivum/Info/Annotation; see Acknowledgements). We firstly, filtered the SNPs removing those not showing allelic changes characteristic of EMS treatment (leaving 89% of the SNPs). Secondly, we selected only those SNPs with a mapping quality of >= 50. Thirdly, we annotated homozygous mutant SNPs as those having the mutant or alternate allele in >85% of the total sequencing reads i.e. wild-type (WT) allele in <=15% of the reads, the remaining SNPs were annotated as heterozygous. Finally, we removed SNPs where the mutant alternate allele was observed at less than 4X coverage and removed SNPs that were not assigned to one of the 21 wheat chromosomes. This resulted in a list of high confidence SNPs for CO calling. COs were called as previously, however, instead of looking for Parent 1/Parent 2 shifts we recorded Homozygous/Heterozygous shifts between windows.

### Gene Conversion calls (GC-Phenotype)

The RefSeq V1.0 anchored SNPs from the 35K array that were used for CO calling were used here to identify short shifts between parents that we use as a potential indicator for GC. Here, no windows or collapsing was used and each change in the encountered parent specific allele from one SNP to the next across the genome was recorded as an event. Here, we assume a correct local reference order and consider each genotype shift even if they are short in length or multiple changes occur in close proximity. Therefore, here we consider both GC and CO events together due to our limited ability to detect the difference via the specific length of the event with only on average 4335 SNPs available for analysis per population with a genome size of 16Gb. This methodology has the potential to include false-positives led by incorrect genome order and incorrect SNP calls. As such, here we rely on the recent advances in the wheat reference sequence assembly and genome ordering alongside the high confidence SNP calls generated by the 35K array. For direct estimates of the number of GC events per RIL we subtract the number of COs from the GC/CO estimate.

To call GCs from exome capture data from the Cadenza TILLING population; SNPs were identified in the data and identified as Homozygous/Heterozygous as previously detailed. GCs (again alongside COs) were called as previously however, instead of looking for Parent 1/Parent 2 shifts we recorded Homozygous/Heterozygous shifts across the genome. Since we observed high variation in the number of SNPs that were available for each EMS treated line (1,700-9,018) alongside a linear relationship between the number of SNPs available and the number of GCs identified (R^2^=0.91595), we normalized GC estimates to reflect a SNP count of 5,000 per line.

### Determining if COs show bias to genic regions

To determine if COs are more or less likely to target genic regions, we compared the number of genes at the CO sites to the number of genes in intervals of the same size across the genome, which contain an array SNP, independently of whether they contained a CO or not. Focusing on only those intervals containing array SNPs eliminates bias from the array SNP positions already being focused in genic regions.

### Cluster analysis

The dendrograms in Figure 1e and 1f were produced using the R package pvclust average linkage method with correlation-based dissimilarity matrix and AU (Approximately Unbiased) p-values were computed by multiscale bootstrap resampling (bootstrap number of 1,000).

### Phylogenetic Analysis of the ***RecQ* Gene Family**

RecQ-like genes were identified by detecting the presence of the helicase conserved C-terminal domain in the proteomes of ten species using HMMER3.1b2 HMMSEARCH (hmmer.org). The inputs to HMMSEARCH were the Pfam hidden Markov model (HMM), Helicase_C (PF00271), and the protein data sets from the following genome annotations: Arabidopsis (Araport11), Medicago (Phytozome, V10), Brachypodium (Phytozome, V12), rice (MSU_RGAP_v7.0), maize (Phytozome, V10), barley high and low confidence genes (IBSC consortium, Mascher et al, 2017), wheat TGACv1 (Clavijo et al 2017), wheat RefSeq v1.0 high and low confidence genes (IWGSC consortium), Marchantia (Marchantia.info), moss (Phytozome V10) and yeast (Ensembl Fungi Release 40). The Helicase_C protein sequences detected were aligned back to the HMM using HMMER3.1b2 HMMALIGN. Gap columns in the alignment were removed and sequences with less than 70% coverage across the alignment were removed to reduce false placement in the tree of sequences with insufficient coverage across the domain. The longest sequence for each gene out of the available set of splice versions was used for phylogenetic analysis. Phylogenetic analysis was carried out using the MPI version of RAxML v8.2.9 (Stamatakis et al 2014) with the following method parameters set: -f a, -x 12345, -p 12345, -# 100, -m PROTCATJTT. The tree was mid-point rooted and visualized using the Interactive Tree of Life (iToL) tool (Letunic and Bork, 2016) to identify the clade containing the RECQ-like homologs. Full length sequences for these homologs were aligned to build a tree containing Arabidopsis, rice and wheat proteins (Figure 4d) and a tree containing proteins from all 10 species (Supplemental Figure S8a). Each alignment was made with PRANK (Löytynoja and Goldman, 2008), then all the columns between the conserved DEAD and RECQ domains were extracted using JalView (Waterhouse et al, 2009) and the alignment was used to build the final tree for the RecQ-like proteins with the above RAxML command.

### Selecting samples for skim sequencing

RILs were selected from the Paragon x Chinese Spring population due to us having whole genome reference sequences available for both accessions. RILs were selected to cover a profile of low, medium and high CO and GC frequency compared to the averages for the population (∼52 for COs and ∼388 for COs and GCs). Low was defined as >= 10% less than the average (CO range 0-46; CO/GC range 0-350), medium was defined as within +/-10% of the average (CO range 47-57; CO/GC range 351-428) and high was defined as >= 10% more than the average (CO range 58-93; CO/GC range 428-540).

### Genomic DNA isolation for skim sequencing

Leaf tissue from each line was ground in liquid nitrogen with a mortar and pestle. Genomic DNA was extracted from the ground tissue using a DNeasy plant mini kit (Qiagen) according to the manufacturer’s protocol (version March 2018) with two alterations. The incubation time with buffer AP1 and RNase was extended to one hour and a new 1.5ml microcentrifuge tube was used for the second elution to prevent dilution of the DNA. For all library preparations the first elution of each sample was used. The genomic DNA was assessed by spectrophotometry using a NanoDrop 2000 (Thermo Fisher) for contamination. DNA concentration was determined using a Quant-iT High Sensitivity double stranded DNA assay kit (Invitrogen) and a Infinite F200 Pro microplate reader (Tecan). DNA integrity was assessed using genomic DNA ScreenTape (Agilent) and a TapeStation 2200 (Agilent).

### Library preparation for skim sequencing

The twelve genomic DNA samples were sheared to 300 bp using a S2 Covaris ultrasonicator (2 cycles 60s, 10% duty factor, intensity of 5, and 200 cycles/burst). A whole genome library was produced for each sample using the KAPA High Throughput library preparation kit (Roche). The standard protocol was followed (Version 5.16), with the modifications listed below. The safe stopping point in the standard protocol at A-tailing was included. For adapter ligation, 3µl of 10µM SeqCap adapters (Roche) were used. The volume of water per reaction was adjusted so the total volume remained 50µl. The dual size selection ratios were adjusted to 0.5x and 0.7x to account for the larger fragment size. Two cycles of PCR were used to amplify the libraries in order to keep PCR duplicates down. The libraries were quantified by Qubit High Sensitivity double stranded DNA assays (Invitrogen) and a Qubit 2 (Invitrogen). Final library yield was between 80ng-200ng. Fragment size was determined by running the libraries on High Sensitivity DNA Bioanalyzer (Agilent) chips. Exact molar concentrations were determined prior to sequencing using the universal KAPA Illumina Library Quantification kit (Roche) on an Applied Biosystems StepOne Plus system.

### Skim sequencing of whole genome libraries

Sequencing was performed on the Illumina MiSeq and NovaSeq6000 platforms. The libraries were pooled equi-molarly for the MiSeq according to the concentrations determined by qPCR. A MiSeq Nano v2 150bp paired-end run was used to quality check the libraries and optimise library pooling for the NovaSeq run. The NovaSeq pooled library was balanced according to the MiSeq read data. The library was run on two S2 150bp paired-end NovaSeq6000 lanes.

### Identification of Paragon-Chinese Spring specific SNPs for validation of skim sequencing

In this analysis we used the Earlham Institute’s Paragon whole genome assembly (at http://opendata.earlham.ac.uk/opendata/data/Triticum_aestivum/EI/v1.1/). From this sequence we simulated 2×150 bp paired end sequencing reads with no errors at 20X coverage across the genome using the short-read simulator dwgsim v0.1.11. To avoid the pitfalls in GC detection noted by Qi et al. (2014) and Wijnker et al. (2013) we aligned these reads back to the Chinese Spring RefSeq V1.0 using BWA-MEM v0.7.10 (Li and Durbin 2009), took only reads aligned in a proper pair (correct orientation and mapped distance) and removed any non-uniquely mapped reads (mapping quality <= 10) using SAMtools (Li et al. 2009). We aligned 1,923,658,991 simulated reads initially and after filtering 1,817,505,942 reads remained resulting in coverage of 11,390,273,387bp of the 14Gb reference genome size at a minimum of 10X. We then called SNPs between Paragon and the Chinese Spring reference using GATK (McKenna et al. 2010). We focused only on homozygous SNPs (alternate allele frequency >80%) and removed SNPs with a quality score less than 30, a depth less than 3 and removed SNPs if 3 or more were defined within a 10bp window. These SNPs are sites where we can accurately discriminate Chinese Spring and Paragon and we refer to this SNP list as “Paragon-Chinese Spring specific SNPs”.

### Identification of Paragon-Chinese Spring COs and GCs from skim sequencing

Here we aligned sequencing reads for each of the 12 lines that were sequenced to the Chinese Spring RefSeq V1.0 using the same methodology as for the simulated Paragon reads detailed above however, with the following additions; we included a duplicate read removal step using Picard tools v2.1.1 and this time for SNP calling using GATK we removed SNPs with a quality score less than 20. Since we are aligning skim sequencing from Paragon x Chinese Spring crosses to a Chinese Spring genome we expect that the alternate SNP alleles we define in the sequencing data will be Paragon specific and therefore these can be used to define COs an GCs. However, our defined SNPs are typically seen at low coverage and as such require validation. We validated our SNPs in the skim sequencing data by comparing the defined alleles to the “Paragon-Chinese Spring specific SNPs” that we previously defined. For each skim sequenced line we cycle through the list of “Paragon-Chinese Spring specific SNPs” i.e. all possible differences between Chinese Spring and Paragon, and for each SNP position; if the skim sequenced line also has a SNP with an alternate allele defined in >80% of the sequencing reads at this position that matches the Paragon allele we define this as Paragon-specific; if the line has a SNP with an alternate allele in <80% of the reads or that does not match Paragon the position is undetermined; if the line has no SNP allele defined at this position we check for mapping coverage in the region (if none the position is undetermined) and if sufficient coverage is observed but no SNP called we check that the Chinese Spring reference allele is found at this position and define this as Chinese Spring specific. We then have a validated list of positions across each of the skim sequenced lines from which we can define COs and GCs where we either define Paragon specific sequence, Chinese Spring specific sequence or have an undetermined call which is removed from the analysis. Due to working with the F8 generation and not F2 as per previous studies we focus on homozygous SNPs only rather than homozygous-heterozygous shifts that are more likely to be mistakenly identified due to incorrectly aligned reads as highlighted by Qi et al. (2014).

In this analysis we defined COs and GCs in sequencing data as the intersection of unbroken runs of markers from a single parent that are surrounded or followed by runs of markers from the other parent. We defined the lengths of the unbroken runs and used this to classify events as GCs in the category of 2bp-10Kbp (while also subsetting these GCs into regions of 20bp-2Kbp and 2-19bp) and classified COs as regions of sequence exchange between parents of >10Kbp (while also subsetting these COs into regions of 10-500Kbp and >500Kbp). Due to the large genome size of wheat it is unknown if GCs/COs may present as longer shifts than were seen in *Arabidopsis* therefore we categorized blocks of; 20bp-2Kbp, 2-10Kbp, 10-500Kbp, >500Kbp and also using blocks of >20Mbp/>40Mbps to allow comparison with our previous array-based analysis. To ensure confident calls for CO/GC determination, we only considered a break in the run of markers from one parent to another if 3 or more SNPs on the run showed such a shift between Paragon and Chinese Spring. Since our resolution of SNPs that discriminate Chinese Spring and Paragon is 1 SNP per ∼540bp we should still be able to effectively detect COs and the majority of GCs using this methodology while eliminating false positive calls.

### Validation of COs and GCs from array analysis with skim sequencing

Using the array, our refinement of a CO is limited to the intersection of two 20Mbp windows. In order to validate CO’s that were defined using the array with COs that were defined using skim sequencing, we looked at the interval 20Mbp each side of each array defined CO. We then determined if this region overlapped a sequence exchange site or GC that we defined from the skim sequencing with high confidence i.e. the region between the SNPs that show the Chinese Spring-Paragon shift. We expected our COs to overlap these shifts that were associated with GCs of >20Mbp. Table 2 details the percentage of COs that overlapped and the average length of the GC that was overlapped was in line with our expectations.

To validate GCs that were defined using the array, we looked at the interval 3Mbp around our array defined GC (representative of the typical genome space between array SNPs if 5000 are analyzed across a 16Gb genome) and determined if this fell within a GC that we defined from the skim sequencing. Table 2 details the percentage of array GCs that overlapped a skim sequencing GC and those overlapping skim sequencing GCs with the same SNP allele were classified as high confidence GCs. As an additional validation for the skim sequenced sample 0004, we identified paired end sequencing reads that spanned Chinese Spring-Paragon GC shift sites and where the SNPs used to define the shifts were close enough in proximity to allow a single read pair to provide coverage for both the Chinese Spring and Paragon SNPs. At 169 shift sites we gained a minimum of 3X coverage for both the Chinese Spring and Paragon SNPs that was made up from single read pairs and this allowed to us calculate a likely false positive rate from the perspective of read misalignment or structural reference related problems. For correlation of array defined GCs with skim sequencing defined GCs we normalized our skim sequencing defined GC frequencies according to the number of SNP sites that show sequencing coverage in the individual datasets where Paragon and Chinese Spring can be discriminated since this has a direct impact on GC calls.

### QTL analysis

Our QTL analysis implemented genetic maps for each individual population that have been previously generated and are publicly available (see Data Availability). QTL calculation and plotting of logarithm of odds (LOD) scores were conducted using R package ‘qtl’, in the first step as a single QTL model employing the extended Haley-Knott method on genotypes. Significant thresholds for the QTLs were calculated from the data distribution. Final QTL LOD scores and effects were received from a multiple QTL model, using the QTL detected in the initial scan.

### Immunolocalisation

Immunolocalisation was performed using the protocol from Higgins et al (2013) using primary antibodies anti-ASY1 (guinea pig) and anti-γH2A.X (rabbit) (Millipore). Secondary antibodies: anti-guinea pig 488 (false coloured to blue for contrast with red) and anti-rabbit alexa 594 (Invitrogen). Images were captured using Nikon NIS-Elements software and processed with the Mexican Hat function. Individual γH2A.X foci co-localising with ASY1 were marked using the counting tool.

## Declarations

### Acknowledgements

DNA sequence was generated by the Earlham Institute-Genomic Pipelines (United Kingdom). We thank Robert King, Christian Schudoma, Cristobal Uauy, and Ksenia Krasileva for providing SNP calls and early access to these SNP calls for the Cadenza TILLING population overlaid onto the IWGSC RefSeq V1. We thank Simon Orford for his assistance obtaining the Paragon x Chinese Spring seeds at the John Innes Centre Germplasm Resources Unit. Assemblies of the Paragon cultivar were generated in the BBSRC funded Strategic LOLA project and we thank Mike Bevan and Bernardo Clavijo for authorizing and facilitating early access to this dataset.

### Authors’ contributions

Array data, genetic maps and QTL methodology were provided by LW. PB identified knockouts for candidate genes from the TILLING population and generated phylogenetic trees (Fig 4d, Fig S8). RJ generated simulated reads for Paragon, performed mapping alignment for skim sequencing datasets, generated figure 2f, performed plant growth. TB performed Illumina sequencing library preparation for skim sequencing samples. JW resolved Chinese Spring-Paragon synteny for candidate genes. JH provided guidance and conceptual support and performed immunolocalization experiments (Fig 2d-e, Fig S6). LG performed; SNP calling in skim sequencing data and definition of parent specific SNPs, methodology development and calls for CO/GC determination in both the array and skim sequencing data, CO/GC landscape analysis, QTL analysis, candidate gene search and manuscript writing/figure preparation. The project was designed, planned and conducted by AH and LG with assistance and expertise from BC, NH, SG and JH. The paper was written by LG and AH with assistance from NH and JH. All authors approved the final manuscript.

### Competing interests

The author(s) declare that they have no competing interests.

### Ethics approval

All plants used in this study were grown in controlled growth chambers complying with Norwich Research Park guidelines. Plant material was supplied from the Germplasm Resources Unit at the John Innes Centre, Norwich, UK.

### Funding

This project was supported by the BBSRC via an ERA-CAPS grant BB/N005104/1, BB/N005155/1 (L.G, A.H), IWYP project grant BB/N020871/1 (R.J), BBSRC funded Strategic LOLA project BB/N002628/1 (JDH) and BBSRC Designing Future Wheat BB/P016855/1 (A.H, L.G). Assemblies of the Paragon cultivar were generated in the BBSRC funded Strategic LOLA project (BB/J003557/1).

### Data availability

Genetic maps and 35K array data for the 13 populations under analysis are publicly available (https://data.cimmyt.org/dataset.xhtml?persistentId=hdl:11529/10996, http://www.cerealsdb.uk.net/cerealgenomics/CerealsDB/axiom_download.php and http://wisplandracepillar.jic.ac.uk/results/JIC_DFW_Watkins_ParWat_Ax_iS_maps.xlsx). The SNPs from the Cadenza TILLING lines are available publicly (https://plants.ensembl.org/Triticum_aestivum/Info/Annotation). The skim sequencing datasets are available (study PRJEB28231) from the European Nucleotide Archive (http://www.ebi.ac.uk/ena/data/view/).

### List of Supplemental materials

Supplemental Notes S1-S9

Supplemental Figures S1-S9

Supplemental Tables S1-S7

