## Supplemental_Information for "Analysis of the recombination landscape of hexaploid bread wheat reveals genes controlling recombination and gene conversion frequency"

#### Supplemental Notes

Supplemental note S1-Supplemental note S7

#### Supplemental Figures

Supplemental Figure S1-Supplemental Figure S9

#### Supplemental Tables

Supplemental Table S1-Supplemental Table S7

#### Supplemental Notes

**Supplemental Note S1:** For the Holliday junction ATP-dependent DNA helicase *RuvB-like* that was associated with CO frequency, we identified two lines with likely knockouts via introduced stop codons and a further six with missense mutations (Supplemental Table S6). We defined CO frequency for each of the eight mutant lines using SNPs that were defined from the TILLING population exome capture data (CO-Phenotype, Methods). To enable comparison, we identified ten control lines from the TILLING population with no mutations in our genes of interest and calculated their CO frequencies (CO-Phenotype). After comparison, the CO frequencies of the knockout *RuvB* lines and control group largely overlapped with average frequencies of 57.4 and 57.6 respectively (Two tailed t test, P=0.9562, t=0.0558, df=16) (Figure 4a, Methods).

**Supplemental Note S2:** For *RecQ-7* that was associated with GC frequency, we identified four lines with likely knockouts via introduced stop codons (Supplemental Table S6) and defined GC frequency for each of the four mutant lines (GC-Phenotype, Methods). We used the same control lines for comparison as used for the *RuvB* analysis but this time calculated their GC frequencies for the lines.

**Supplemental Note S3:** The average number of SNPs that were available to perform our GC phenotyping analysis across the Cadenza TILLING lines under analysis was 5,207 with a range from 1,700-9,018. Following on from this 462.5 was the average number of CO/GCs that were identified across the TILLING lines with 405.5 likely GCs. Previously, using the array SNPs, with on average 4335 SNPs available for analysis per population, we defined an average of only 104 GCs per RIL across the 13 populations (Supplemental Table S2). This increase in GC detection for the TILLING lines is thought to be due to a combination of the increased number of SNPs for analysis alongside the larger population size of the TILLING population (1,200 lines compared to the average size of 158 RILs for the 13 previously analyzed populations). This is supported by our identification of 335.5 GCs per RIL in the Paragon x Chinese Spring population where we have a larger number of 8,369 SNPs available and a larger population size than many of the the other RIL populations at 269.

**Supplemental Note S4:** From Figure 4d, we were able to identify the three homoeologous wheat homologs of the *Arabidopsis* recombination candidate genes *RecQ4A* and *RecQ4B*. We then used the Cadenza TILLING population, to ascertain if knockouts of these homoeologous genes showed CO frequency phenotypes. We were able to identify 18 knockouts across the three homoeologs (Supplemental Table S6). Knockouts of our gene candidates resulted in a decrease in the average GC frequency per line from 498.5 in the control group to 413.7, 460.7 and 446.9 for sub-genome A, B and D respectively (GC-Phenotype), although only the decrease from knockouts of sub-genome A was statistically significant (Two tailed t test, sub-genome A; P=0.0163, t=2.6833, df=16, sub-genome B; P=0.3710, t=0.9243, df=14, sub-genome D; P=0.2533, t=1.2466, df=7). Looking at CO frequency, knockouts of our gene candidates resulted in a decrease in the average CO frequency per line from 59 in the control group to 53.5, 53.5 and 54.8 for sub-genome A, B and D respectively (CO-Phenotype), although none of the decreases were statistically significant (Two tailed t test, sub-genome A; P=0.1240, t=1.6236, df=16, sub-genome B; P=0.2426, t=1.2200, df=14, sub-genome D; P=0.3610, t=0.9496, df=12).

**Supplemental Note S5:** From Figure 4d, we were able to identify the homoeologous B and D sub-genome copies of our candidate gene *RecQ-7* on chromosome 2A. There was an additional low confidence gene that was also closely related to our homoeologous trio and observed on chromosome 2D. We then used the Cadenza TILLING population, to ascertain if knockouts of these homoeologous genes showed similar phenotypes that were observed with the knockout on chromosome 2A. We were able to identify three knockouts of the homoeolog on chromosome 2B, one knockout of the homoeolog on chromosome 2D and an additional knockout of the closely associated low confidence gene on chromosome 2D (Supplemental Table S6). Knockouts of our gene candidates resulted in a decrease in the average GC frequency per line from 498.5 in the control group to 414.4 and 449.4 for sub-genomes B and D respectively (GC-Phenotype). Although a decrease was observed (Supplemental Figure S9), it was not statistically significant, potentially due to the low number of knockouts under analysis (Two tailed t test, sub-genome B; P=0.1504, t=1.5457, df=11, sub-genome D; P=0.095, t=1.8656, df=9).

**Supplemental Note S6:** For GC frequency (as per GC-Phenotype, Methods) we identified multiple robust QTL that were seen in populations other than the Paragon x Chinese spring population. Firstly, we identified ATP-dependent DNA helicase *PIF2* from the Paragon x CIMMYT 47 analysis, for this gene we were unable to identify any Cadenza TILLING lines showing knockouts. We were able to identify two TILLING lines showing missense mutations with sift scores <0.05 in this gene, however, these mutants showed an average GC frequency of 486.5 which, although lower than the control group (498.5), was not significantly different (Welch two tailed t test, p=0.8973, t=0.1628, df=1) (Supplemental Table S7).
Secondly, we identified a gene encoding the protein HIRA from the Paragon x Watkins 94 analysis. For this gene we were able to identify six TILLING lines with potential knockouts i.e. stop codons gained (Supplemental Table S7). These mutants showed an average GC frequency of 439.4 which, although lower than the control group (498.5) by almost 60 GCs, was not significantly different (Welch two tailed t test, p=0.1632, t=1.4945, df=11).
Finally, we identified the WPP domain-interacting protein 1 from the Paragon x Baj analysis. For this gene we were unable to identify any Cadenza TILLING lines showing knockouts. However, we were able to identify seven TILLING lines with missense mutations with sift scores <0.05 (Supplemental Table S7). These mutants showed an average GC frequency of 454.5 which, although again lower than the control group (498.5), was not significantly different (Welch two tailed t test, p=0.2069, t=1.3235, df=14). Interestingly, when we went on to define CO frequency for these mutants to determine if our GC-Phenotype could translate to CO frequency, we observed a significant decrease in CO frequency in the mutant group compared to the control group (Welch two tailed t test, P=0.0277, t=2.4562, df=14) with average CO frequencies of 50.7 and 59 for mutant and control respectively. Therefore, this gene could be an important candidate for further analysis.

**Supplemental Note S7:** The candidate gene for our third QTL encoded the protein HIRA on chromosome 4B that is needed for chromatin reassembly during DSB repair (Li, X. and Tyler, J.). HIRA mediates DNA synthesis-independent nucleosome assembly and in mammalian cells has been shown to take over for nucleosome assembly on newly replicated DNA in the absence of CAF-1 (Brachet et al., 2015). While, CAF-1 interacts in a DNA damage-dependent manner with the ATP-dependent RNA helicase RecQ (Bernstein et al., 2014; Hoek et al., 2011). This

QTL therefore may highlight, within the Paragon x Watkins 94 population, the use of an alternative but complementary pathway to the previously defined QTLs for recombination.

Our final QTL identified the WPP domain-interacting protein 1 on chromosome 5A as a candidate gene alongside the Regulator of chromosome condensation (RCC1) gene. WPP domain-interacting protein 1 mediates and enhances nuclear envelope docking of RANGAP proteins; the cycling of Ran between its GTP- and GDP-bound forms is catalysed by (RCC1) and RANGAP resulting in an intracellular concentration gradient of RanGTP that acts as a GPS for cells with regard to meiotic spindle formation that can affect chromosome segregation (Zhao, et al., 2008; Kalab and Heald, 2008; Cesario and McKim, 2011). Furthermore, Cadenza TILLING lines showing missense mutations for this gene showed a drop in GC frequency that translated to a significant drop in CO frequency. It is evident from all our QTL analyses, that the maximum peak in our defined QTL interval is consistently within 1Mbp of a functionally relevant gene that has previously been linked to either homologous recombination or DSB repair and these genes represent potential targets for wheat breeding programs in the drive to increase recombination rate.

#### Supplemental Figures

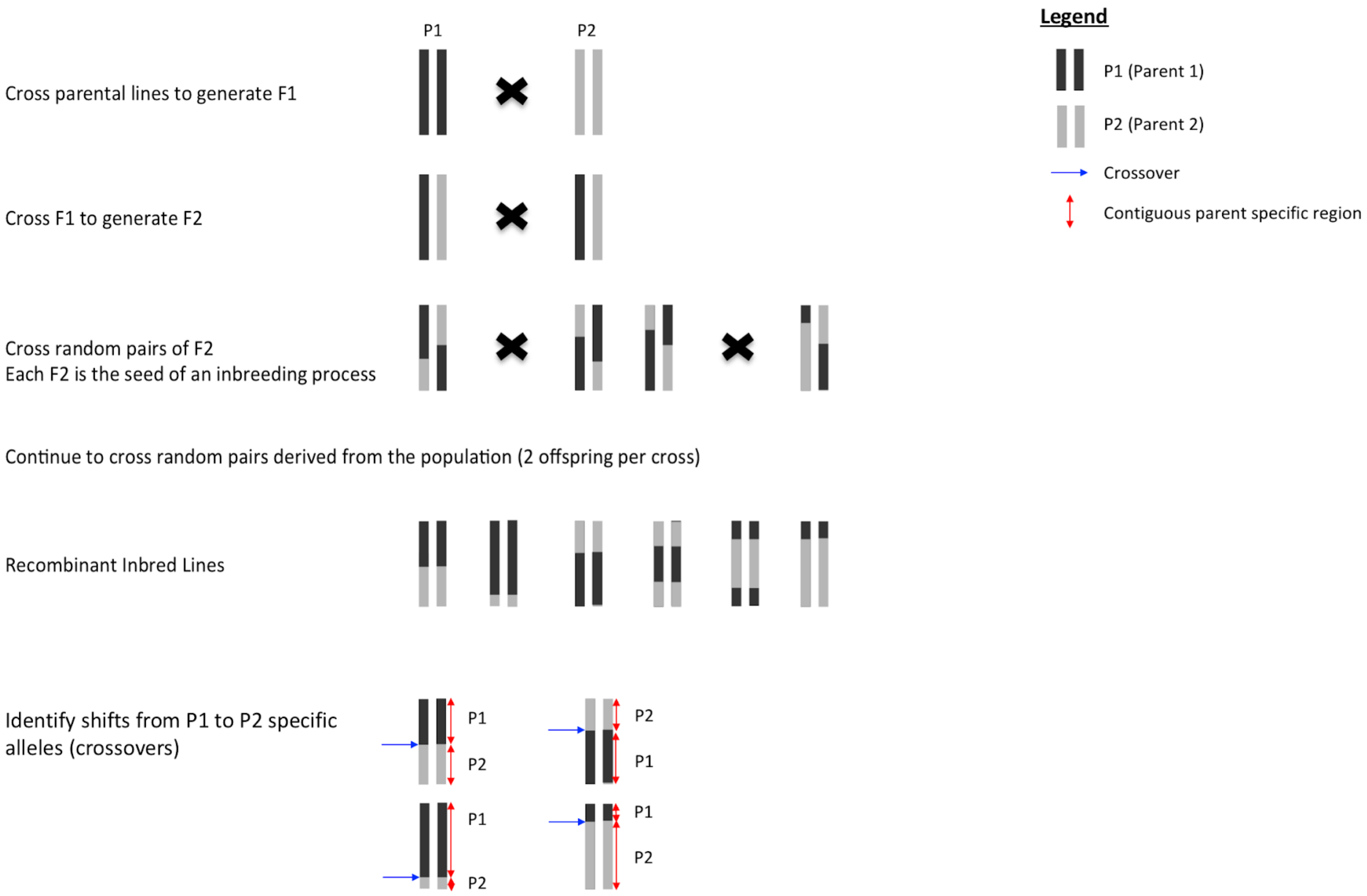


**Supplemental Figure S1. Construction of a RIL line and CO identification.** Example of the construction of a Recombinant inbred line (RIL) where a set of diploid chromosomes represents an individual and each parental genotype is represented by either black or grey. Each individual resultant RIL is a mosaic of the parental lines with widespread homozygosity between chromosome pairs. By comparing the genotype of each RIL to that of its parental founders, it is possible to see shifts in genotype from alleles specific to Parent 1 to alleles specific to Parent 2. These shifts can be used as an estimate of recombination points or chromosomal COs across the genome to predict where they occurred during population generation.

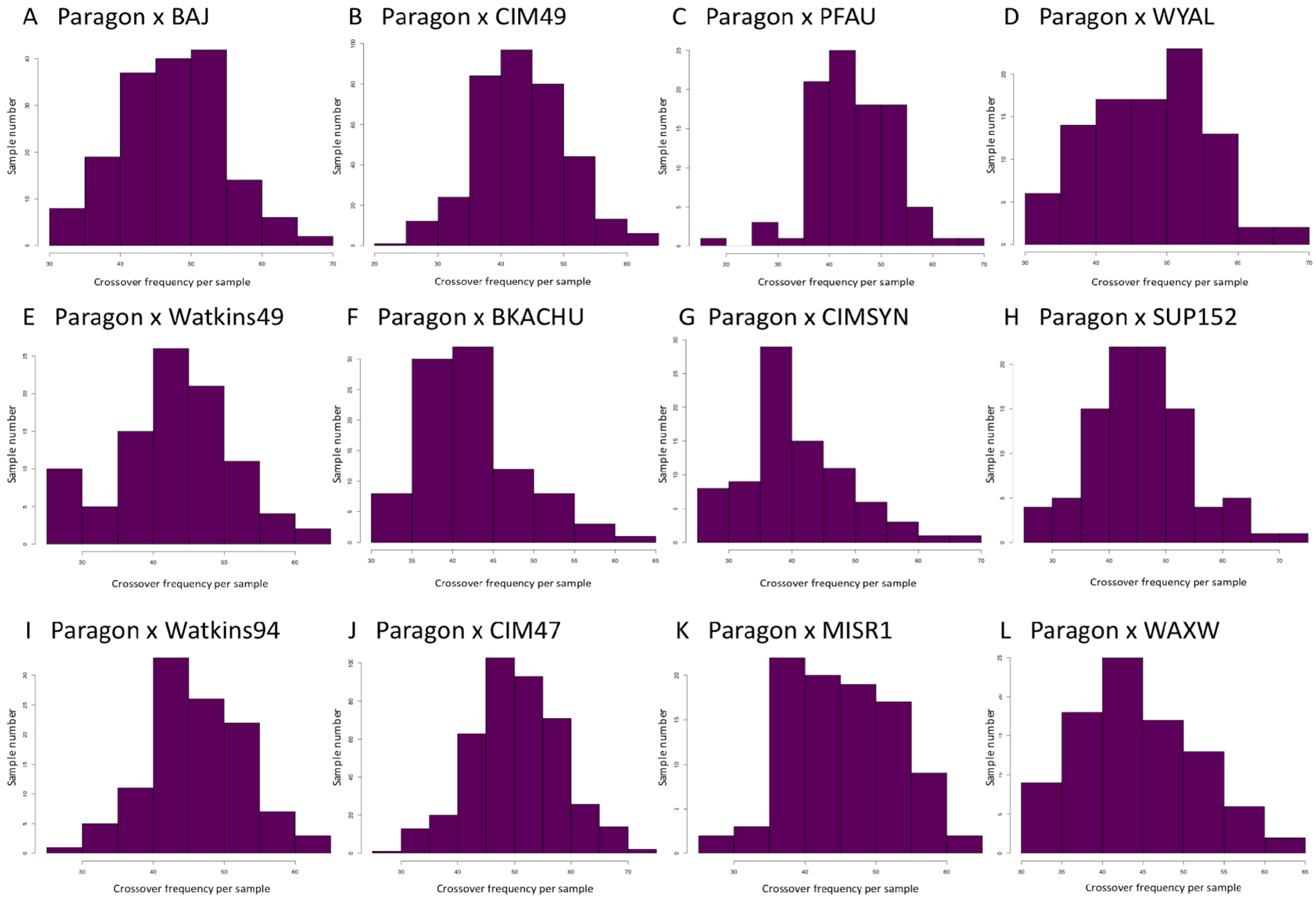


**Supplemental Figure S2. Frequency of COs per RIL.** For each of the populations under analysis figures A-L represent the number of COs recorded for each RIL (CO frequency per sample) as a frequency histogram.

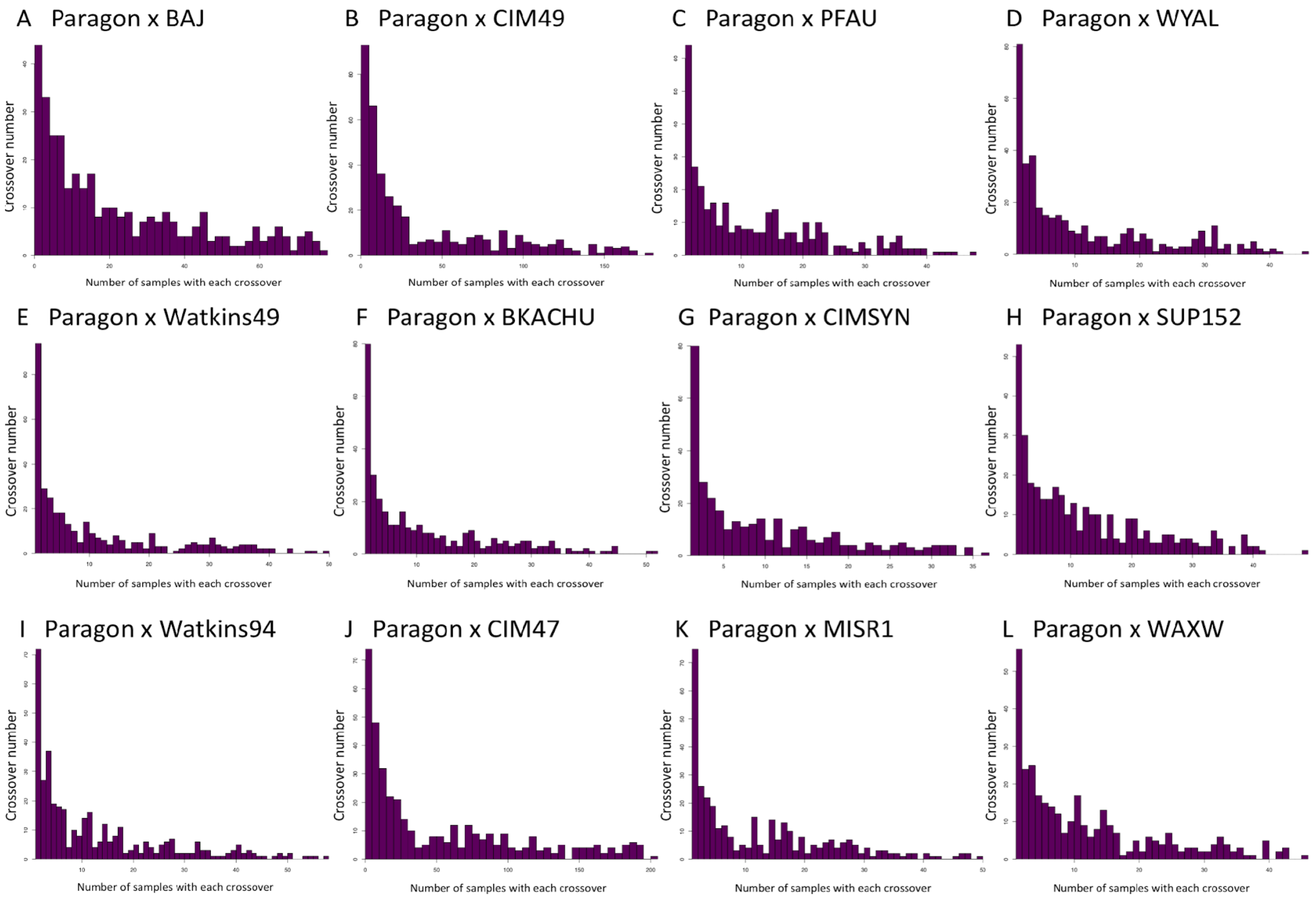


**Supplemental Figure S3. Number of shared COs per population.** For each of the populations under analysis figures A-L represent the number of RILs sharing each recorded CO (Number of samples with each CO) as a frequency histogram.

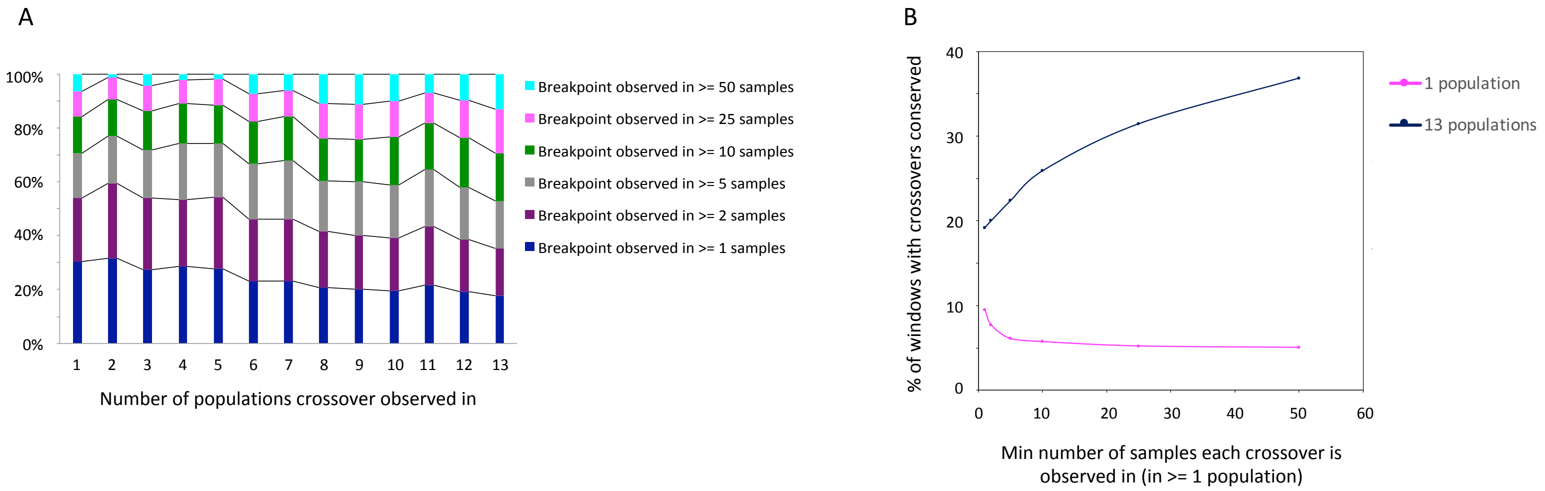


**Supplemental Figure S4. CO conservation between populations.** (A) Stacked bar chart summing all observed COs in each category (number of populations from 1-13). Bar charts are sub divided into coloured blocks representing how many RILs shared the CO within a population (see legend). (B) Scatter Plot displaying the minimum number of RILs each CO has to have been observed in (x axis) versus the percentage of COs that are conserved (y axis) across the relevant number of populations (1 and 13 respectively).

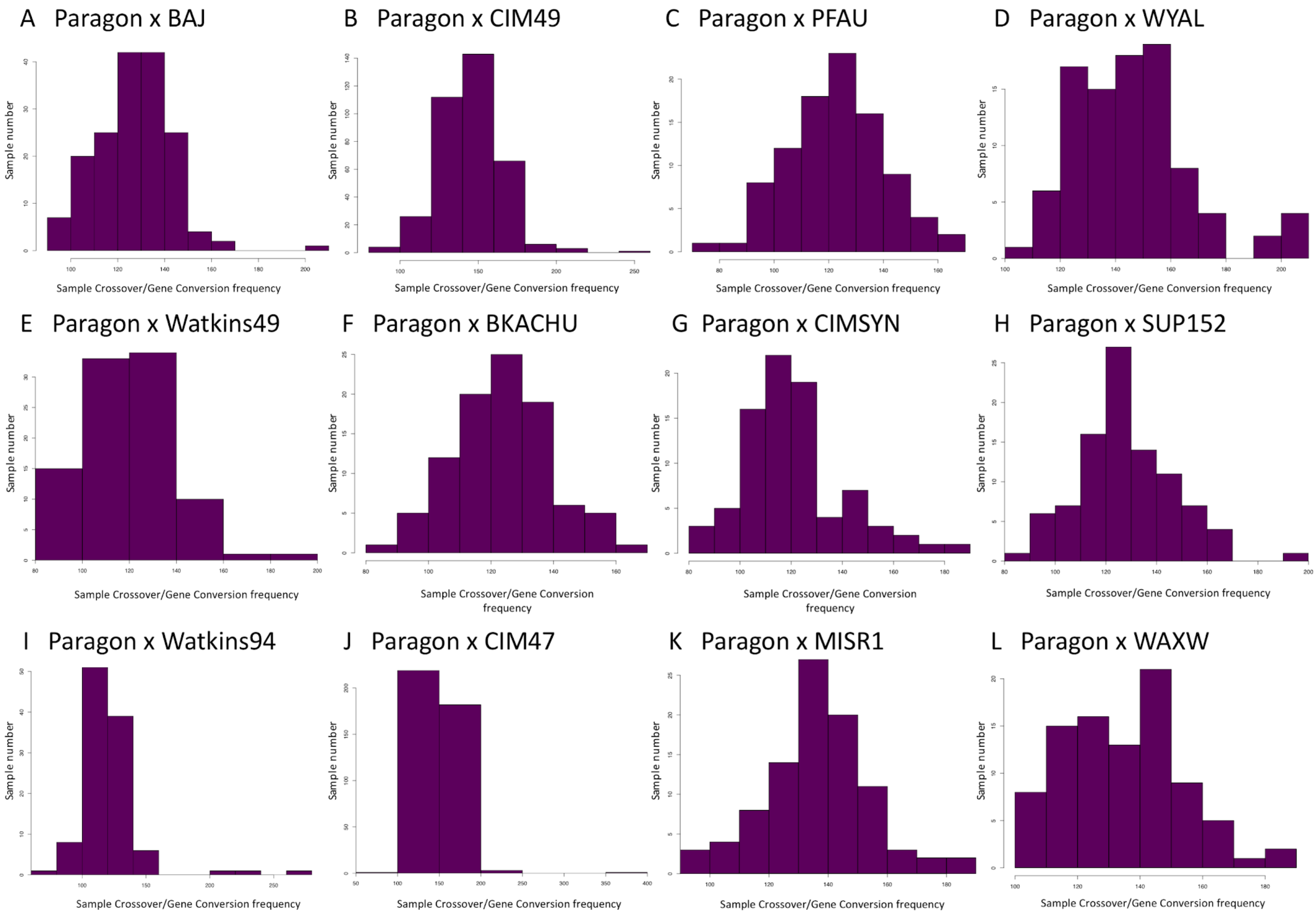


**Supplemental Figure S5. Frequency of COs and GCs per RIL.** For each of the populations under analysis figures A-L represent the number of COs and GCs i.e. total sequence exchange events recorded for each RIL (CO frequency per sample) as a frequency histogram.

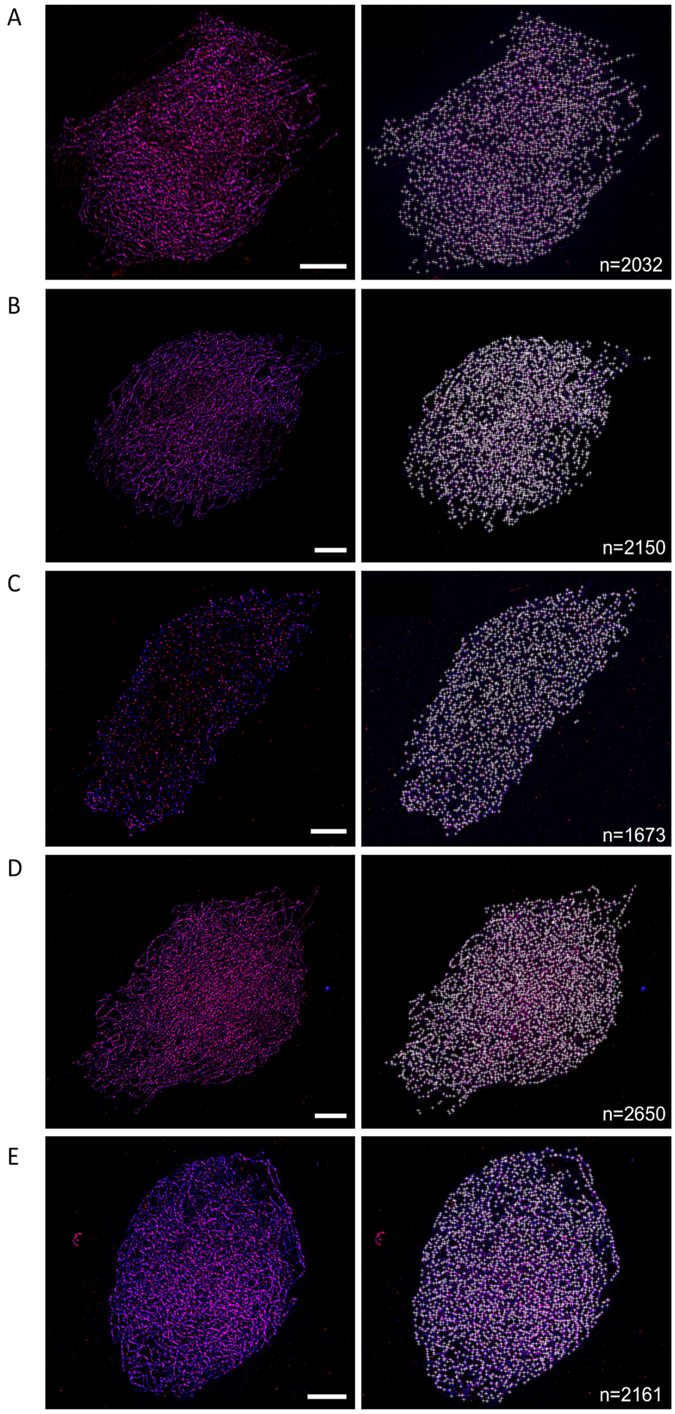


**Supplemental Figure S6. Defining DSBs in hexaploid wheat.** Immunolocalisation of the chromosome axis protein ASY1 (blue) and γH2A.X (red) a marker for DNA double strand breaks on hexaploid wheat leptotene male meiotic nuclei. (A-E) five individual replicate nuclei with the image to the left the original nuclei and the rightmost image the original nuclei with γH2A.X foci marked that co-localise with ASY1. Scale bar = 10μM. Mean γH2A.X foci = 2133 (n=5), SD = 313, SE = 157.

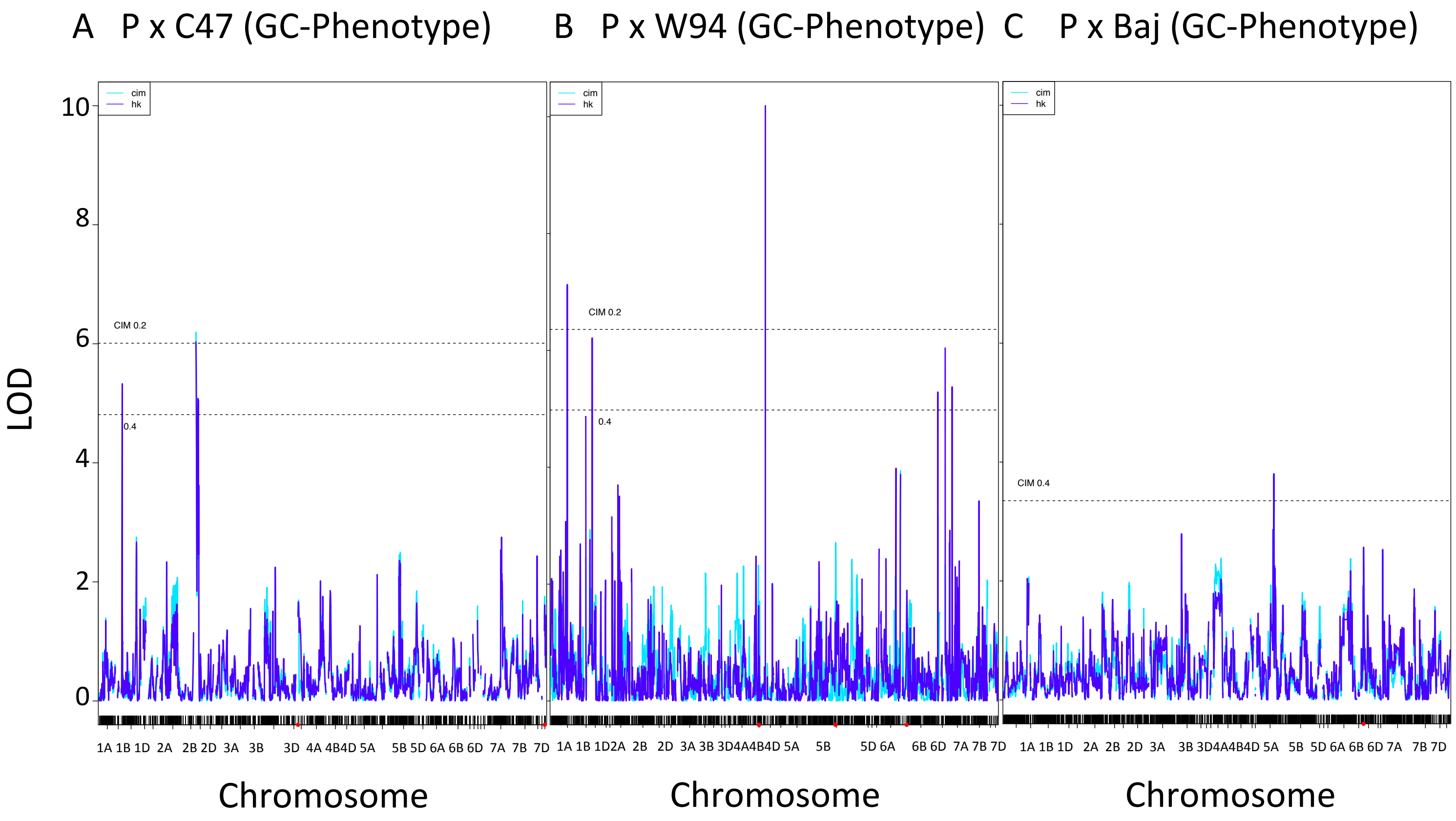


**Supplemental Figure S7. Output from additional QTL analysis.** QTL analysis output for the additional three populations that yielded significant associations for GC-Phenotype ($p<0.05$). Detailing LOD scores plotted over the respective linkage groups i.e. chromosomes.

A

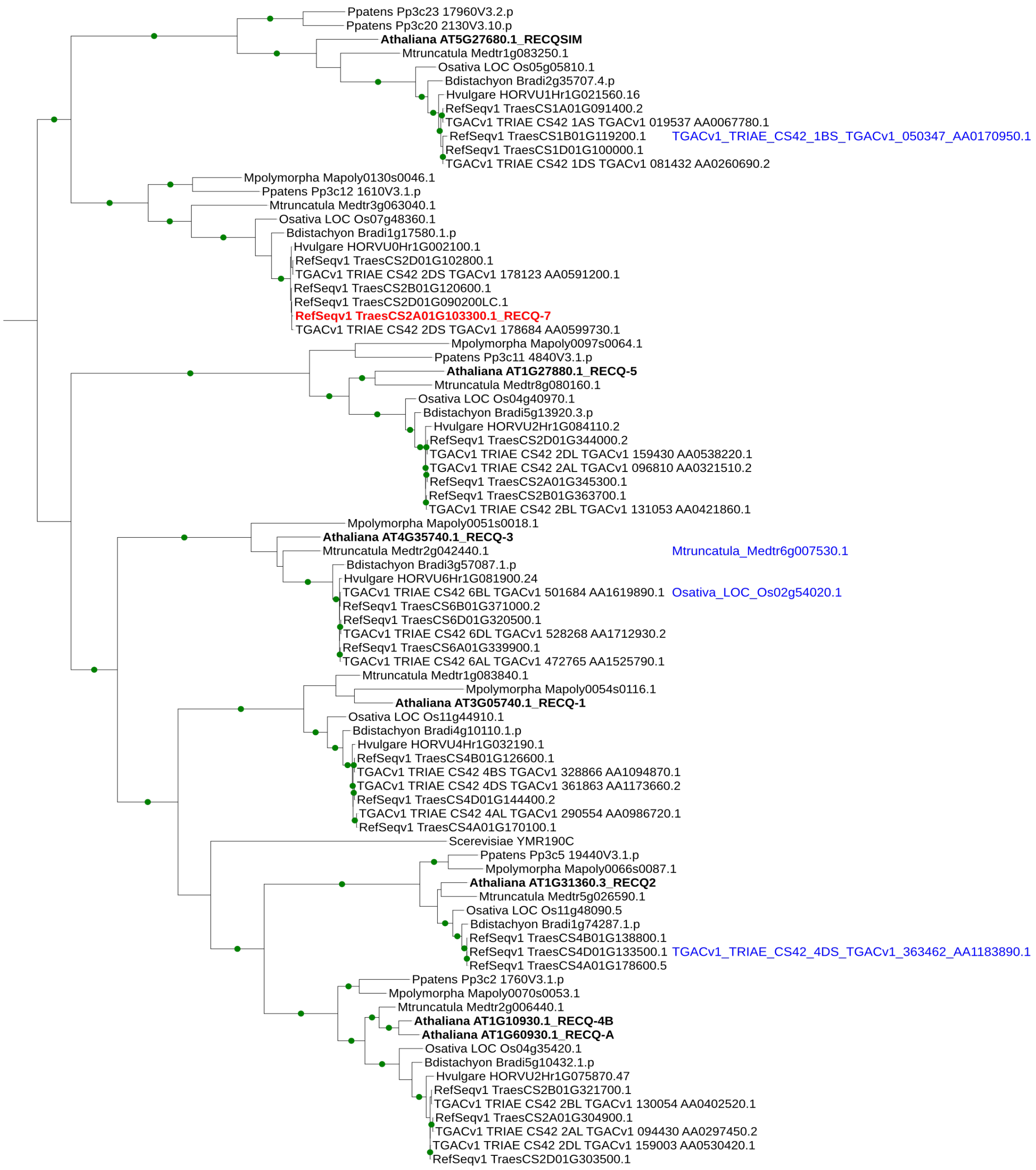


**Supplemental Figure S8. Phylogenetic Analysis of the *RecQ* Gene family. (A)** Phylogenetic tree of all *RecQ*-like genes across ten species with sequence similarity to the RecQ helicase family (Methods). Bootstrap values >= 90 % are shown as green dots on the branches. Additional gene identifiers shown in blue are truncated genes belonging to the RecQ family at the position indicated, based on a BLASTP search of the Helicase_C family.

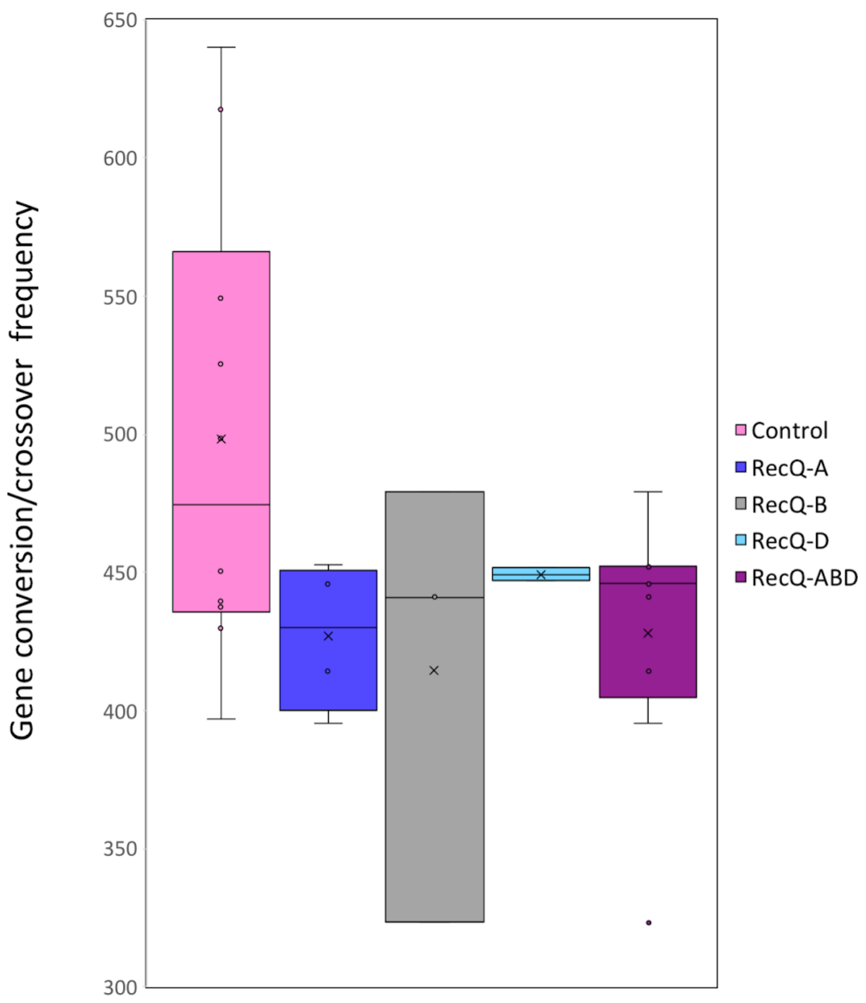


**Supplemental Figure S9. Examination of homoeologs of candidate genes from QTL analysis.** Box plot comparison of the knockout *RecQ-7* lines with the control lines, defining CO/GC frequency using GC-Phenotype. Knockouts are shown for the candidate *RecQ-7* gene on chromosome 2A (RecQ-A) and its predicted homoeologs on chromosome 2B (RecQ-B) and 2D (RecQ-D). The RecQ-D group also includes knockouts for the low confidence gene on chromosome 2D that also shows high similarity to the homoelogous gene trio. Finally, all knockouts, independently of which homoeologous gene is knocked out, are pooled into a single group for comparison (RecQ-ABD).

### Supplemental Tables

**Table S1. Detailing the parental founders of the 13 populations that were used in this study.**

| Parent 1 name | Parent 1 information | Parent 2 name | Parent 2 information |
|---|---|---|---|
| Paragon | UK elite line | Chinese Spring | Landrace (reference variety) |
| Paragon | UK elite line | Watkins 1190049 | Watkins Bread wheat landrace diversity collection |
| Paragon | UK elite line | Watkins 1190094 | Watkins Bread wheat landrace diversity collection |
| Paragon | UK elite line | BAJ | CIMMYT Breeders line |
| Paragon | UK elite line | BKACHU | CIMMYT Breeders line |
| Paragon | UK elite line | CIMMYT 47 | CIMMYT Breeders line |
| Paragon | UK elite line | CIMMYT 49 | CIMMYT Breeders line |
| Paragon | UK elite line | CIMMYT SYN | CIMMYT Breeders line |
| Paragon | UK elite line | MISR1 | CIMMYT Breeders line |
| Paragon | UK elite line | PFAU | CIMMYT Breeders line |
| Paragon | UK elite line | SUP152 | CIMMYT Breeders line |
| Paragon | UK elite line | WAXW | CIMMYT Breeders line |
| Paragon | UK elite line | WYAL | CIMMYT Breeders line |

**Table S2. Population statistics for the 13 populations under investigation.** Detailing for each of the 13 populations; population size, the number of 35K array SNPs that were consistently assayed across the population and informative i.e. polymorphic between Parent 1 and Parent 2, the number of unique COs identified per sample, the average number of COs per sample and the average number of GCs per sample.

| **Parent 1 name** | **Parent 2 name** | **Population Size** | **Number of 35K SNPs assayed** | **Unique COs in population** | **COs per sample** | **GCs per sample** |
|---|---|---|---|---|---|---|
| Paragon | Chinese Spring | 269 | 8369 | 466 | 51.9 | 335.5 |
| Paragon | Watkins 1190049 | 94 | 4010 | 357 | 43.5 | 75.9 |
| Paragon | Watkins 1190094 | 108 | 4057 | 383 | 46.7 | 73.4 |
| Paragon | BAJ | 168 | 3765 | 354 | 47.8 | 79.1 |
| Paragon | BKACHU | 94 | 3884 | 343 | 42.7 | 80.4 |
| Paragon | CIMMYT 47 | 406 | 4789 | 394 | 50.9 | 97.9 |
| Paragon | CIMMYT 49 | 361 | 4502 | 407 | 43.8 | 101.7 |
| Paragon | CIMMYT SYN | 83 | 4021 | 332 | 40.8 | 79.9 |
| Paragon | MISR1 | 94 | 4391 | 340 | 45.6 | 89.2 |
| Paragon | PFAU | 94 | 4629 | 342 | 45.5 | 76.7 |
| Paragon | SUP152 | 94 | 4075 | 345 | 45.9 | 81.1 |
| Paragon | WAXW | 90 | 4085 | 329 | 45.1 | 88.4 |
| Paragon | WYAL | 94 | 4618 | 389 | 48.1 | 96.6 |

**Table S3. Shared COs within the 13 populations under investigation.** Detailing for each of the 13 populations; the percentage of the total number of COs that are shared between various numbers of RILs.

| Sample Number | % COs shared (466 total) PxCS | % COs shared (357 total) PxWat49 | % COs shared (383 total) PxWat94 | % COs shared (354 total) PxBAJ | % COs shared (343 total) PxBKACHU | % COs shared (394 total) PxCIM47 | % COs shared (407 total) PxCIM49 | % COs shared (332 total) PxCIMSYN | % COs shared (340 total) PxMISR1 | % COs shared (342 total) PxPFAU | % COs shared (345 total) PxSUP152 | % COs shared (329 total) PxWAXW | % COs shared after bp collapse (389 total) PxWYAL |
|---|---|---|---|---|---|---|---|---|---|---|---|---|---|
| 1 | 6.65 | 15.4 | 10.4 | 7.1 | 14.6 | 3.8 | 6.1 | 12.3 | 12.4 | 11.7 | 9.0 | 12.2 | 10.8 |
| 2 | 6.44 | 10.9 | 8.4 | 5.4 | 8.7 | 4.8 | 5.2 | 11.7 | 9.7 | 7.0 | 6.4 | 4.9 | 10.0 |
| 3 | 7.51 | 8.1 | 7.0 | 5.4 | 8.7 | 2.5 | 3.4 | 8.4 | 7.6 | 7.9 | 8.7 | 7.3 | 9.0 |
| 4 | 5.58 | 7.0 | 9.7 | 4.0 | 6.1 | 3.6 | 3.7 | 6.6 | 6.5 | 6.1 | 5.2 | 7.6 | 9.8 |
| >=2 | 93.3 | 84.6 | 89.6 | 92.9 | 85.4 | 96.2 | 93.9 | 87.7 | 87.6 | 88.0 | 91.0 | 87.8 | 89.2 |
| >=5 | 73.8 | 58.5 | 64.5 | 78.2 | 61.8 | 85.3 | 81.6 | 60.8 | 63.8 | 67.0 | 70.7 | 68.1 | 60.4 |
| >=10 | 58.6 | 40.6 | 46.7 | 61.3 | 43.1 | 70.6 | 62.9 | 41.9 | 48.2 | 48.8 | 48.4 | 48.3 | 41.1 |
| >=20 | 43.6 | 22.1 | 24.0 | 43.5 | 23.3 | 57.4 | 46.9 | 16.3 | 25.3 | 24.0 | 24.6 | 23.4 | 22.6 |
| >=30 | 34.1 | 12.9 | 13.1 | 31.6 | 9.9 | 46.4 | 36.6 | 5.7 | 9.4 | 10.8 | 9.9 | 10.9 | 12.1 |
| >=40 | 29.0 | 2.5 | 7.0 | 21.5 | 2.9 | 43.1 | 33.7 | 0 | 2.9 | 2.0 | 1.7 | 3.3 | 1.3 |
| >=50 | 24.5 | 0.3 | 1.8 | 13.6 | 0.6 | 39.6 | 30.7 | 0 | 0.3 | 0 | 0 | 0 | 0 |
| >=100 | 6.0 | 0 | 0 | 0 | 0 | 19.3 | 13.0 | 0 | 0 | 0 | 0 | 0 | 0 |
| >=125 | 2.6 | 0 | 0 | 0 | 0 | 13.5 | 6.6 | 0 | 0 | 0 | 0 | 0 | 0 |
| >=150 | 0 | 0 | 0 | 0 | 0 | 9.6 | 3.7 | 0 | 0 | 0 | 0 | 0 | 0 |
| >=200 | 0 | 0 | 0 | 0 | 0 | 0.3 | 0 | 0 | 0 | 0 | 0 | 0 | 0 |

**Table S4. Shared COs between the 13 populations under investigation.** Detailing the percentage of the total number of COs that are shared between 1-13 populations according to how many RILs within specific populations have the CO. Cumulative percentage totals for each row equal 100%.

| In >=1 popln the min no of samples brkpt is in | % windows with brkpts in 1 popln | % windows with brkpts in 2 poplns | % windows with brkpts in 3 poplns | % windows with brkpts in 4 poplns | % windows with brkpts in 5 poplns | % windows with brkpts in 6 poplns | % windows with brkpts in 7 poplns | % windows with brkpts in 8 poplns | % windows with brkpts in 9 poplns | % windows with brkpts in 10 poplns | % windows with brkpts in 11 poplns | % windows with brkpts in 12 poplns | % windows with brkpts in 13 poplns |
|---|---|---|---|---|---|---|---|---|---|---|---|---|---|
| 1 | 9.5394 | 7.7302 | 6.5788 | 4.7697 | 4.7697 | 6.25 | 5.09868 | 3.4539 | 7.0723 | 7.40131 | 7.23684 | 10.8552 | 19.243 |
| 2 | 7.7186 | 7.0325 | 6.6895 | 4.2881 | 4.8027 | 6.51801 | 5.31732 | 3.6020 | 7.3756 | 7.71869 | 7.54716 | 11.3207 | 20.068 |
| 5 | 6.1302 | 4.9808 | 4.9808 | 4.0229 | 4.0229 | 6.5134 | 5.55555 | 3.6398 | 8.2375 | 8.62068 | 8.23754 | 12.6436 | 22.413 |
| 10 | 5.7649 | 4.43 | 4.656 | 3.3259 | 3.3259 | 5.76496 | 4.8780 | 3.547 | 7.5388 | 9.09090 | 7.76053 | 13.9689 | 25.942 |
| 25 | 5.2478 | 3.4985 | 4.0816 | 2.6239 | 2.9154 | 4.95626 | 3.79008 | 3.7900 | 8.1632 | 8.74635 | 6.70553 | 13.994 | 31.487 |
| 50 | 5.0847 | 0.8474 | 2.5423 | 0.8474 | 0.8474 | 5.08474 | 3.38983 | 4.6610 | 10.169 | 9.74576 | 5.9322 | 13.9830 | 36.864 |

**Table S5. Samples selected for whole genome skim sequencing.** Detailing the samples sequenced from the Paragon x Chinese Spring population, the number of COs defined for each sample and the number of GCs defined for each sample (those classed as high are in bold type and low are underlined; see Methods for details). Also detailing the number of sequencing reads aligned to the IWGSC RefSeqV1 Chinese Spring reference genome plus average depth and percentage of genome mapped. Finally, the number of SNPs called are detailed alongside the number of Paragon-Chinese Spring parent specific sites that have sequence information in the skim sequenced samples.

| RIL Name (Paragon x Chinese Spring population) | CO frequency (35K array) | GC frequency (35K array) | Aligned sequencing reads-post filtering | Average depth of coverage | % of reference genome mapped | Number of SNPs identified | Number of Paragon-CS specific sites with information (31,327,143 total) |
|---|---|---|---|---|---|---|---|
| 0055 | **73** | **429** | 617,259,125 | 6.74 | 92.21 | 13,984,026 | 29,182,483 |
| 0334 | **70** | **507** | 577,017,395 | 6.33 | 91.21 | 19,739,698 | 27,012,789 |
| 0200 | 54 | **461** | 632,964,899 | 6.91 | 91.76 | 19,285,179 | 28,830,075 |
| 0270 | 54 | **483** | 623,216,599 | 6.73 | 92.05 | 14,937,252 | 28,278,916 |
| 0008 | **65** | 413 | 553,203,920 | 6.16 | 90.65 | 26,542,822 | 27,609,483 |
| 0119 | **69** | 409 | 602,689,927 | 6.65 | 91.43 | 20,542,948 | 27,779,818 |
| 0005 | 55 | 363 | 581,127,334 | 6.39 | 90.91 | 23,687,781 | 27,926,026 |
| 0018 | 52 | 386 | 496,591,667 | 5.48 | 90.91 | 17,775,994 | 27,712,617 |
| 0097 | 53 | 322 | 651,920,209 | 7.20 | 90.69 | 29,376,821 | 28,166,225 |
| 0004 | 34 | 378 | 483,449,595 | 5.37 | 90.40 | 18,188,963 | 27,408,941 |
| 0010 | 37 | 319 | 559,978,269 | 6.22 | 90.39 | 21,867,247 | 27,831,695 |
| 0281 | 39 | 306 | 629,503,914 | 6.82 | 92.36 | 14,797,624 | 29,149,453 |

**Table S6. Validation of CO frequency QTLs from Paragon x Chinese Spring population using the Cadenza TILLING population.** Detailing; candidate gene, length of CDS, TILLING line with likely effector mutation in candidate gene, consequence of mutation (stop codon or missense mutation), from exome sequencing of line the coverage of the reference with wild type reads (WT) and mutant allele containing reads (Mut).

| Candidate gene | Length of CDS (bp) | Annotation | TILLING Line | Category | Mutation Consequence | WT Coverage | Mut Coverage | CDS Position |
|---|---|---|---|---|---|---|---|---|
| TraesCS2A01G103300 | 2679 | Chromosome 2A ATP-dependent helicase RecQ (*RecQ-7)* | Cadenza0927 | het5hom3 | stop_gained | 28 | 11 | 228 |
| | | | Cadenza1716 | het5hom3 | stop_gained | 0 | 18 | 940 |
| | | | Cadenza0560 | het5hom3 | stop_gained | 14 | 21 | 1318 |
| | | | Cadenza1279 | het5hom3 | stop_gained | 6 | 8 | 1758 |
| TraesCS2B01G120600 | 2607 | Chromosome 2B *RecQ-7* B homoeolog | Cadenza1733 | het5hom3 | stop_gained | 17 | 12 | 130 |
| | | | Cadenza0393 | het5hom3 | stop_gained | 19 | 11 | 1307 |
| | | | Cadenza1675 | het5hom3 | stop_gained | 0 | 33 | 1887 |
| TraesCS2D01G102800 | 2682 | Chromosome 2D *RecQ-7* D homoeolog | Cadenza0207 | - | stop_gained | 0 | 8 | 205 |
| TraesCS2D01G090200 | 306 | Chromosome 2D *RecQ-7* D homoeolog | Cadenza0284 | - | stop_gained | 13 | 13 | 172 |
| TraesCS6A01G026300 | 2112 | Chromosome 6A Holliday junction ATP-dependent DNA helicase RuvB | Cadenza1616 | het5hom3 | stop_gained | 0 | 2 | 289 |
| | | | Cadenza1495 | het5hom3 | stop_gained | 17 | 18 | 787 |
| | | | Cadenza1530 | het5hom3 | Splice_acceptor_variant | 0 | 4 | |
| | | | Cadenza2095 | het5hom3 | missense_variant | 34 | 4 | 1162 |
| | | | Cadenza0129 | het5hom3 | missense_variant | 7 | 6 | 82 |
| | | | Cadenza1709 | het5hom3 | missense_variant | 0 | 35 | 443 |
| | | | Cadenza0962 | het5hom3 | missense_variant | 0 | 9 | 458 |
| | | | Cadenza1410 | het5hom3 | missense_variant | 67 | 51 | 476 |
| TraesCS2A01G304900 | 3423 | Chromosome 2A *Arabidopsis RecQ4A/4B* homolog | Cadenza1636 | het5hom3 | stop_gained | 0 | 3 | 1577 |
| | | | Cadenza0387 | het5hom3 | stop_gained | 11 | 13 | 1624 |
| | | | Cadenza1371 | het5hom3 | stop_gained | 53 | 48 | 3325 |
| | | | Cadenza0861 | het5hom3 | stop_gained | 14 | 10 | 3363 |
| | | | Cadenza0300 | het5hom3 | stop_gained | 25 | 8 | 257 |

| | | | | | | | | |
|---|---|---|---|---|---|---|---|---|
| | | | Cadenza0213 | het5hom3 | stop_gained | 0 | 11 | 258 |
| | | | Cadenza0297 | het5hom3 | stop_gained | 25 | 12 | 1513 |
| | | | Cadenza0865 | het5hom3 | stop_gained | 0 | 51 | 1585 |
| TraesCS2B01G321700 | 3579 | Chromosome 2B *Arabidopsis RecQ4A/4B* homolog | Cadenza1575 | het5hom3 | stop_gained | 0 | 12 | 463 |
| | | | Cadenza0586 | - | stop_gained | 0 | 57 | 1575 |
| | | | Cadenza1993 | - | stop_gained | 83 | 70 | 1774 |
| | | | Cadenza0250 | - | stop_gained | 0 | 29 | 3286 |
| TraesCS2D01G303500 | 3606 | Chromosome 2D *Arabidopsis RecQ4A/4B* homolog | Cadenza0668 | het5hom3 | stop_gained | 0 | 23 | 148 |
| | | | Cadenza0947 | het3hom2 | stop_gained | 1 | 6 | 258 |
| | | | Cadenza1267 | het5hom3 | stop_gained | 8 | 5 | 1213 |
| | | | Cadenza1178 | - | stop_gained | 0 | 26 | 1761 |
| | | | Cadenza1700 | - | stop_gained | 23 | 5 | 2574 |
| | | | Cadenza0316 | - | stop_gained | 15 | 17 | 3546 |

**Table S7. Validation of CO frequency QTLs from various populations using the Cadenza TILLING population.** Detailing; candidate gene, length of CDS, TILLING line with likely effector mutation in candidate gene, consequence of mutation (stop codon or missense mutation), from exome sequencing of line the coverage of the reference with wild type reads (WT) and mutant allele containing reads (Mut).

| Candidate gene | Length of CDS (bp) | Annotation | TILLING Line | Category | Mutation Consequence | WT Coverage | Mut Coverage | CDS Position |
|---|---|---|---|---|---|---|---|---|
| TraesCS2B01G103300 | - | Chromosome 2B PIF2 | Cadenza1346 | het5hom3 | missense_variant | 8 | 10 | 1049 |
| | | | Cadenza0750 | het2hom2 | missense_variant | 1 | 3 | 914 |
| TraesCS5A01G257000 | 1635 | Chromosome 5A WPP | Cadenza0871 | het5hom3 | missense_variant | 9 | 15 | 763 |
| | | | Cadenza0104 | het5hom3 | missense_variant | 0 | 23 | 142 |
| | | | Cadenza0190 | het5hom3 | missense_variant | 14 | 10 | 625 |
| | | | Cadenza0453 | het5hom3 | missense_variant | 14 | 15 | 616 |
| | | | Cadenza0706 | het5hom3 | missense_variant | 4 | 6 | 592 |
| | | | Cadenza2048 | het5hom3 | missense_variant | 4 | 5 | 532 |
| | | | Cadenza2010 | het5hom3 | missense_variant | 20 | 4 | 497 |
| TraesCS4B01G006700 | 2994 | Chromosome 4B HIRA | Cadenza1457 | - | stop_gained | 0 | 39 | 175 |
| | | | Cadenza0142 | - | stop_gained | 16 | 16 | 553 |
| | | | Cadenza0627 | - | stop_gained | 3 | 6 | 553 |
| | | | Cadenza1024 | - | stop_gained | 0 | 21 | 582 |
| | | | Cadenza1720 | - | stop_gained | 4 | 5 | 910 |
| | | | Cadenza0406 | - | stop_gained | 0 | 19 | 2035 |
